# Single cell analysis of neonatal naïve CD8α^+^ T cells reveals novel subsets bridging the innate-adaptive spectrum

**DOI:** 10.64898/2026.03.05.706693

**Authors:** Adam Geber, Brandon Groff, Jordan McMurry, Nathan Laniewski, Adam Tyrlik, Connor Kean, Ruoqiao Wang, Darline Castro-Melendez, Janiret Narvaez-Miranda, Natalie Vance, Gloria Pryhuber, Timothy Mosmann, Brian D. Rudd, Juilee Thakar, David J. Topham, Andrew Grimson, Kristin Scheible

## Abstract

There is growing evidence that neonates harbor innate-like CD8a^+^ T cell subsets that contribute to both protection and hyper-inflammatory states. It remains unclear, however, where these innate-like features are found among the many conventional and unconventional T cell populations that can upregulate the CD8 receptor. Further delineation of these unique populations and functions, with a focus on CD8ab co-expression, will enable studies that seek to understand the unique immune features in conventional T cell populations that are present during fetal and early postnatal life. We used cord blood from infants across the full viable gestational age range to examine phenotypic and transcriptional heterogeneity, with a particular focus on the naïve T cell pool. We report a set of fetally-derived and innate-like naïve CD8αβ^+^ T cells (‘FITs’) that are marked by their KLRG1^+^CD161^+^ phenotype, unique transcriptomic features and which are sparsely detected in adult peripheral blood. Additionally, using T cell receptor repertoire profiling, we can distinguish FITs from well-described and semi-invariant unconventional T cell populations such as mucosa-associated invariant T cells. Our delineation of FITs’ unique features will enable future investigation into their ontogeny and tissue distribution, and ultimately their role in immune-related outcomes in preterm infants.

## Introduction

The human neonatal immune system must balance competing demands to protect and grow the host during the perinatal period. Neonates are particularly susceptible to infection by viruses and encapsulated bacteria that require cellular immune responses for control, yet they must also maintain immune tolerance as they begin harboring diverse commensal microbes at barrier sites. Additionally, preterm neonates have increased susceptibility to diseases of prematurity with hypothesized immunopathological mechanisms including necrotizing enterocolitis and bronchopulmonary dysplasia.^1,2^ Thus, while infants are immunologically naive, their immune systems have a tolerogenic bias that can be overcome or perhaps even bypassed with appropriate and sufficient stimuli to mount an inflammatory response.^3,4^ Despite marked differences between the infant and adult immune system, the majority of infants, including those born prematurely, avoid serious infection, which raises the question of alternative pathways to protection during infancy. The ’layered ontogeny’ model of immune development reconciles these competing demands by describing distinct but overlapping waves of hematopoiesis during fetal development.^5,6^ Hematopoietic stem cells emerging from the fetal liver and bone marrow give rise to a sequence of myeloid and lymphoid cells unique to the developmental stage and with different functions that reflect the needs of the developing embryo.^7^

Some of the earliest T cell populations to arise are the “unconventional” T cells (UTCs), both αβ and γδ chain-expressing. Classically, several qualities distinguish UTCs from conventional naïve αβ T cells. αβ UTCs such as mucosa-associated invariant T cells (MAITs) or type 1 natural killer T cells (iNKTs) express a semi-invariant or restricted TCR and can acquire effector functions and tissue homing capacities prior to thymic egress.^8,9^ They are capable of responding to non-peptide antigens often presented via monomorphic major histocompatibility complex (MHC)-like molecules (e.g. MR1, CD1d, BTN3A1).^10,11 8,9^ Many UTCs also express surface receptors previously associated with innate immune cells and are poised for ’bystander activation’ via TCR-independent and cytokine-mediated signals, particularly IL-12 and IL-18.^8^ In contrast, naïve conventional cells persist after thymic selection in a quiescent but poised state until their peptide antigen-specific and TCR-dependent activation, at which point they are licensed to differentiate and give rise to effector or memory daughter cells in varying proportions.^12^ Finally, UTCs that express the CD8 coreceptor such as MAITs will preferentially display the TCR-dampening CD8αα homodimer, similar to terminally differentiated CD8αα^+^ T cells.^13,14^

Comparisons of UTC populations across the human lifespan have found that umbilical cord blood contains MR1-reactive, iNKT and γδ T cell populations that are unique relative to adult and even postnatal infant circulation, suggesting that UTCs can vary in their TCR usage, coreceptor expression, antigen reactivity and functional capacities depending on their developmental origins.^15–19^ Interestingly, there is also growing evidence from human cohorts and murine models suggesting that even conventional neonatal αβ T cells can possess innate-like features and may even persist into adulthood.^20^ For example, recent studies have shown that neonatal and infant naïve CD8^+^ αβ T cells in circulation are more prone to effector differentiation than their adult counterparts and express a unique set of transcription factors related to innate-like immune functions and/or T cell development.^21–23^ These results parallel prior descriptions of changes in neonatal naïve CD8^+^ αβ T cells’ phenotype and cytokine production capacity that vary linearly with increasing gestational age (GA) at birth.^24–27^ Collectively, these findings indicate that the neonatal T cell compartment contains populations that blur classical boundaries between the innate and adaptive immune systems. However, without simultaneous measure of cellular phenotype, TCR repertoire and gene expression to directly contrast and compare these subpopulations across the conventional-unconventional spectrum, it is unclear if fetally-enriched CD8α^+^ populations represent a unique conventional CD8αβ^+^ αβ T cell subpopulation or rather comprise a subset of the innate-like lymphocytes that predominantly express the CD8αα coreceptor.

We hypothesized that human umbilical cord blood contains naïve CD8αβ^+^ αβ T cell subpopulations with innate-type characteristics bridging conventional and unconventional CD8α^+^ T cell functions. To address this hypothesis, we sought to define a fetally-derived and innate-like CD8αβ^+^ αβ T cell population (‘FITs’) based on a minimum set of criteria including 1) a naïve CD45RA^+^CD45RO^-^CD27^+^ expression pattern, 2) expression of surface phenotypic markers and transcriptome common to unconventional T cell populations and 3) presence in all cord blood samples at greater relative abundance compared to adult peripheral blood. We leveraged cord blood mononuclear cells (CBMC) from neonates of varying gestational age as well as healthy adult peripheral blood mononuclear cells (PBMCs) to profile changes in the composition of the unconventional and CD8α^+^ T cell compartments during late second through third trimester fetal development. Our findings indicate that infant circulating naive CD8α^+^ T cells are heterogeneous, comprising both conventional naïve recent thymic emigrants (RTEs) and are enriched for a subset of naïve fetal innate-like CD8αβ^+^ T cells (FITs) that are functionally and phenotypically distinct from other conventional and unconventional subsets. Distinct functions within the naïve, conventional fetal T cell compartment may provide an alternative, previously unstudied source of either protection or immunopathology in the naïve neonatal host.

## Results

### The neonatal naïve CD8α^+^ compartment is heterogeneous and phenotypically distinct from adult naïve CD8α^+^ T cells

To determine the population structure of circulating naïve CD8a^+^ T cells during late gestation, and to identify phenotypic features that define a naïve fetally-enriched conventional CD8ab^+^ T cell population, we first compared pre-term (32-37 weeks GA, n = 10) and full term (37-40 weeks GA, n = 10) infant CBMC to healthy adult PBMC samples (n = 10) by flow cytometry. Replicate subsamples from each subject underwent *in vitro* stimulation with IL-12/IL-18, PMA/ionomycin, or media alone to determine cells’ cytokine expression under bystander versus mitogen activation conditions. Hierarchical gating confirmed expected distribution of naive (i.e. CD45RA^+^CCR7^±^CD27^+^) and memory subsets of conventional CD8a^+^ T cells, i.e. naïve subsets represented a greater proportion of events in infants compared to adults (Fig 1a). After exclusion of TCRγδ^+^ events, CD56^+^ events and MAITs (CD3^+^TCRVα7.2^+/-^MR1:5-OP-RU^+^), we observed a cord blood enriched naïve CD8αβ-expressing T cell population marked by co-expression of KLRG1 and CD161 (KLRB1) (Fig 1b, 1c). Few KLRG1^+^CD161^+^ naive CD8a^+^ T cells expressed the MAIT invariant α-chain (TCRVα7.2^-^) or exhibited MR1:5-OP-RU tetramer reactivity, indicating that these cells were less likely to be pre-natal, unexpanded MAITs despite their co-expression of the common MAIT surface markers KLRG1 and CD161 (Fig 1c, S1a).

**Figure 1:**
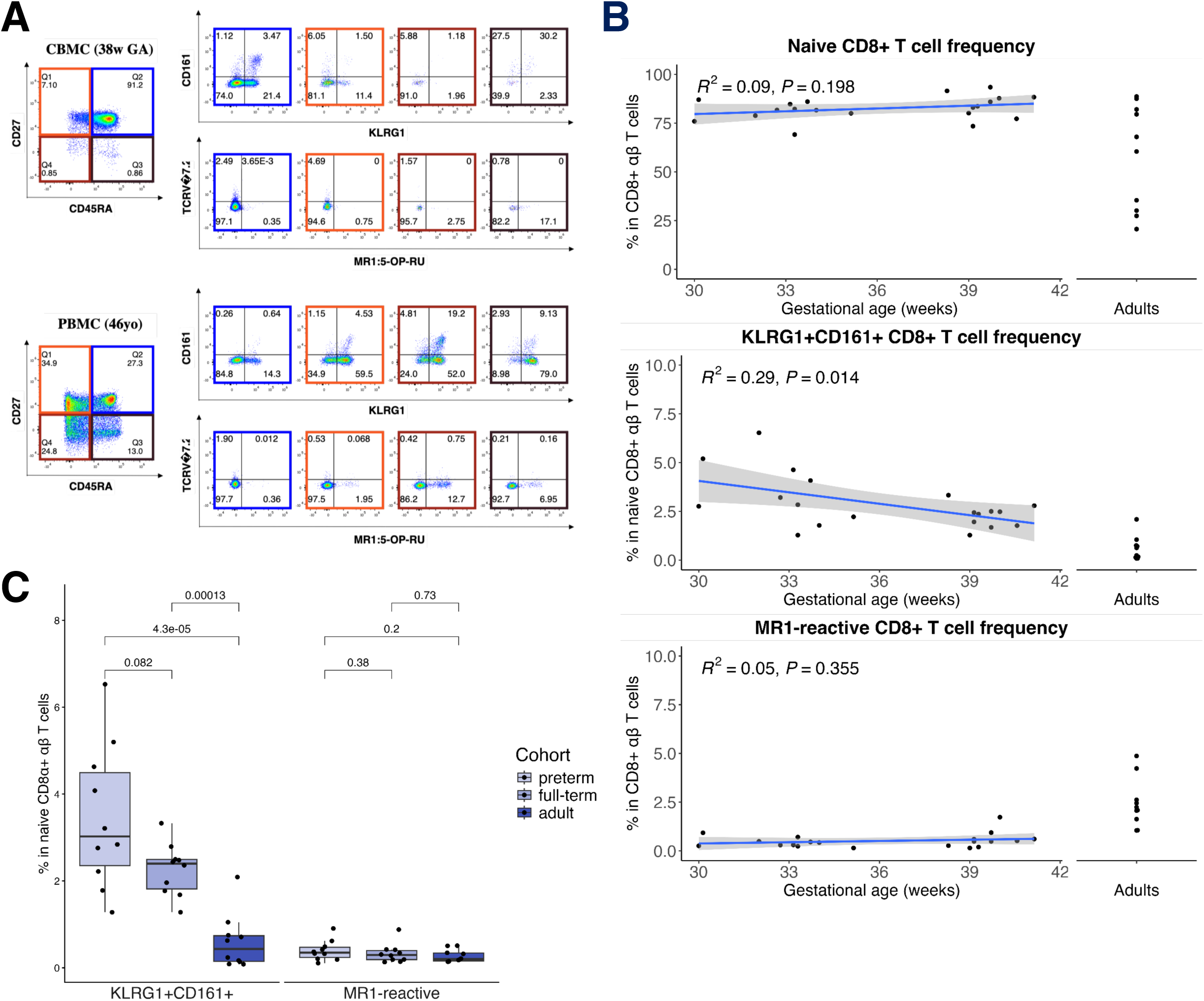
Umbilical cord blood is enriched for a population of naïve CD8α^+^ αβ T cells relative to adult blood. A) Scatterplots showing manually gated CD45RA by CD27 subpopulations for representative infant and adult subjects with inset per-quadrant frequencies. Adjacent scatterplots show the relative staining for KLRG1/CD161 and TCRVα7.2/MR1:5-OP-RU tetramer among the corresponding naïve and memory subpopulations. B) Quantification of the relative frequency of naïve (CD45RA^+^CD27^+^), naïve KLRG1^+^CD161^+^ and MR1-reactive CD8α^+^ αβ T cells among listed infant and adult CD8α^+^ αβ T cells populations. The relationship between population frequency and infant gestational age was assessed using Spearman’s correlation analysis. C) Quantification of the relative frequency of KLRG1^+^CD161^+^ and MR1-reactive populations among naïve CD8α^+^ αβ T cells per cohort. Significance for comparisons of population frequency between age cohorts was determined with the Wilcoxon rank sum test.

To resolve the population structure of the CD8α^+^ T cell compartment among infants and adults in an unbiased fashion, we performed high dimensional clustering of the cellular phenotyping dataset using a modified FlowSOM pipeline (SOMnambulate).^28^ Live singlet lymphocytes were selected from all events and then classified into major immune populations through combinatorial lineage markers (i.e. CD3, CD4, CD8α, CD8β, CD56, TCRγδ). Clusters corresponding to CD8α^+^ T cells (i.e. CD3^+^CD4^-^CD8α^+^CD8β^+/-^CD56^-^TCRγδ^-^) were isolated and reanalyzed using the remaining flow cytometry parameters, which included immunophenotyping markers as well as the intracellular cytokines TNF, IFNγ and IL-8. UMAP dimensionality reduction of these flow cytometry data demonstrated clear separation by CD45RA/RO isoform and CD27/CCR7 expression, recapitulating standard phenotypic distinctions between naive and memory cellular populations (Fig 2a, S2a). After annotating cell clusters according to their memory phenotypic marker expression, we found that two naïve CD8αβ^+^ T cell clusters (12,19) co-expressed KLRG1 and CD161. These clusters also expressed the IL-18 receptor subunit IL18R1 and the transcription factor PLZF (Fig 2b). PLZF is essential for the thymic maturation of UTCs including MAIT, NKT, and certain γδ T cell subsets, such as the phosphoantigen-responsive Vδ2Vγ9 subset.^18,29^ Based on these observations, we assigned clusters 12 and 19 as putative "fetal innate-like T cells" (FITs). FITs could be further distinguished by their divergent expression of the lymphoid homing marker CCR7 and the integrin subunit CD103 (ITGAE), which interacts with epithelial E-cadherin and can be found on subsets of both RTEs and tissue-resident CD8αβ^+^ T cells (Fig 2c). MAITs also co-expressed KLRG1 and CD161, but unlike FITs, they expressed neither CCR7 nor CD27 and were CD45RO^+^. The two CD8a^+^ MAIT clusters (1,9) could be distinguished from FITs by their TCRVα7.2 expression and reactivity to a MR1 tetramer loaded with 5-OP-RU, a strong agonist ligand for MAITs (Fig 2c). Notably, the MAIT population also expressed lower CD8b and higher CD45RO compared to FITs. This phenotypic profile is consistent with the differentiated memory phenotype reported for circulating MAITs.^30^

**Figure 2:**
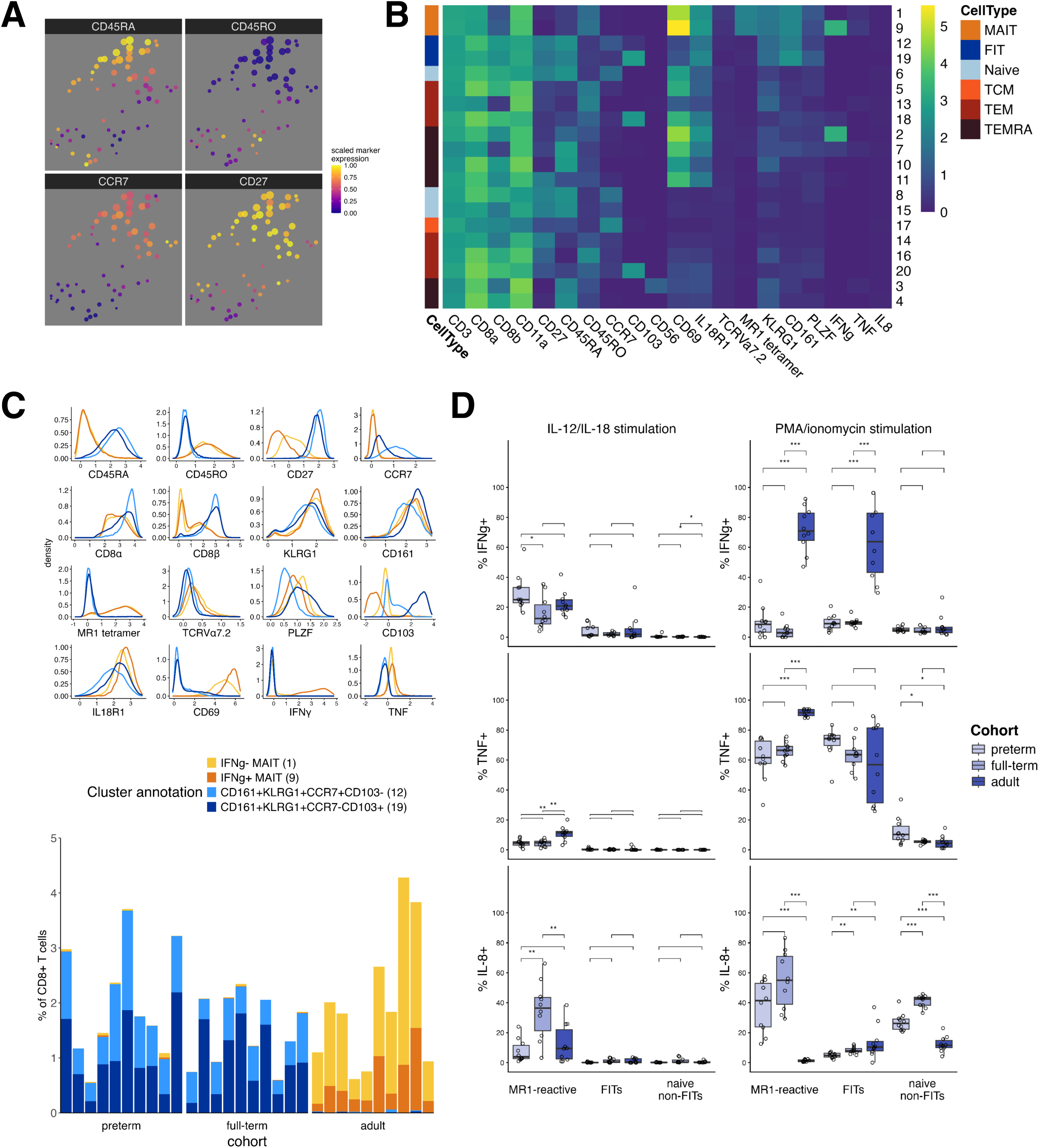
Fetal innate-like T cells are marked by co-expression of KLRG1 and CD161 and produce TNF following PMA/ionomycin stimulation. A) UMAP dimensionality reductions showing expression of specified phenotypic features among FlowSOM-derived nodes. Expression values are scaled from 0 – 1 after trimming extreme events. Data shown in Figures 2A-C correspond to IL-12/IL-18-stimulated cells only. B) Summary heatmap showing median log-scaled expression value for each arcsinh-transformed parameter (columns) across cell clusters (rows, k = 20). Clusters are further annotated by expression of memory phenotypic (i.e. CD45RA/RO, CCR7, CD27) markers. C) Density plots contrasting log-scaled marker expression by MAIT (1/9) and FIT (12/19) clusters (top) with relative frequency of the same clusters among total CD8α^+^ αβ T cells for each subject (bottom). Age cohorts are plotted separately and subjects are ordered along the x-axis according to increasing gestational and biological age. D) Quantification of the relative frequency of cytokine production among MR1-reactive, FIT (KLRG1^+^CD161^+^) and non-FIT (i.e. no co-expression of KLRG1 and CD161) naïve CD8α^+^ αβ T cells following IL-12/IL-18 (left) or PMA/ionomycin (right) stimulation. Subjects are grouped by age cohort as shown. Significance for comparisons of population frequency between age cohorts was determined with the Wilcoxon rank sum test. Nonsignificant pairwise comparisons are not shown.

Despite expression of IL18R1, FITs did not appear to be activated by IL-12/IL-18 treatment in mixed cell culture as determined by their absence of surface expression of CD69 and lack of detectable intracellular IFNγ, TNF or IL-8 under these conditions (Fig 2c, 2d). In contrast, the two MAIT clusters were activated (i.e. CD69^+^) by IL-12/IL-18 treatment but only one cluster produced IFNγ, consistent with previously reported functional heterogeneity among MAITs (Fig 2c).^31^ Subpopulations of MR1-reactive neonatal CD8α^+^ T cells produced IFNγ or IL-8 but not TNF under these stimulation conditions (Fig 2d). Because FITs were minimally responsive to bystander activation and did not produce detectable cytokines under these conditions, we examined the same set of donor cells following PMA/ionomycin stimulation. While stimulated infant naïve CD8α^+^ T cells expressed minimal IFNγ regardless of surface phenotype, most FITs expressed TNF and not IL-8 whereas KLRG1^-^CD161^-^ naïve populations expressed IL-8 with minimal TNF (Fig 2d, S2b). These results indicate that FITs are a major source for the TNF-predominant effector profile known to be enriched in preterm infants, and that they are functionally distinct from the IL-8-predominant CD8^+^ RTE populations extensively described in full term cord blood.^26,32–35^

### Single cell transcriptomics reveals age-based heterogeneity among CD8α^+^ and CD8αβ^+^ lymphoid populations

Flow cytometry revealed a subpopulation of CD8αβ^+^ T cells in cord blood that overlapped phenotypically with adult naïve conventional CD8ab^+^ T cells and also with unconventional T cells, particularly MAITs. However, flow cytometry is limited in its ability to distinguish populations with shared defining markers but distinct functional programs, as may be the case with neonatal naïve CD8αβ^+^ T cells. To identify the transcriptional programs that might further distinguish these populations, we performed CITE-seq on CD8α-expressing T cells and innate lymphoid populations isolated from cord blood samples drawn from gestational ages 22-40 weeks (n=16) and peripheral blood from 4 healthy adult donors. Similar to the experimental conditions used to generate the flow cytometry results described above, replicate wells from each donor’s sample were stimulated with IL-12/IL-18 prior to sorting and sequencing. Stimulated and unstimulated CBMC and PBMC were separately hashed and stained before sorting on the following populations: natural killer (NK) cells (CD3^-^CD56^+^), γδ T cells (CD3^+^TCRγδ^+^), MAITs (TCRVα7.2^+/-^MR1:5-OP-RU^+^), FITs (KLRG1^+^CD161^+^TCRVα7.2^-^MR1:5-OP-RU^-^), and otherwise conventional CD8^+^ T cells (CD3^+^CD4^-^CD8α^+^) (Fig 3a, S3a). Sorted populations from each donor were pooled with target cell counts intended to enrich for rarer populations and to capture the breadth of heterogeneity in antigen and transcript expression within each population.

**Figure 3:**
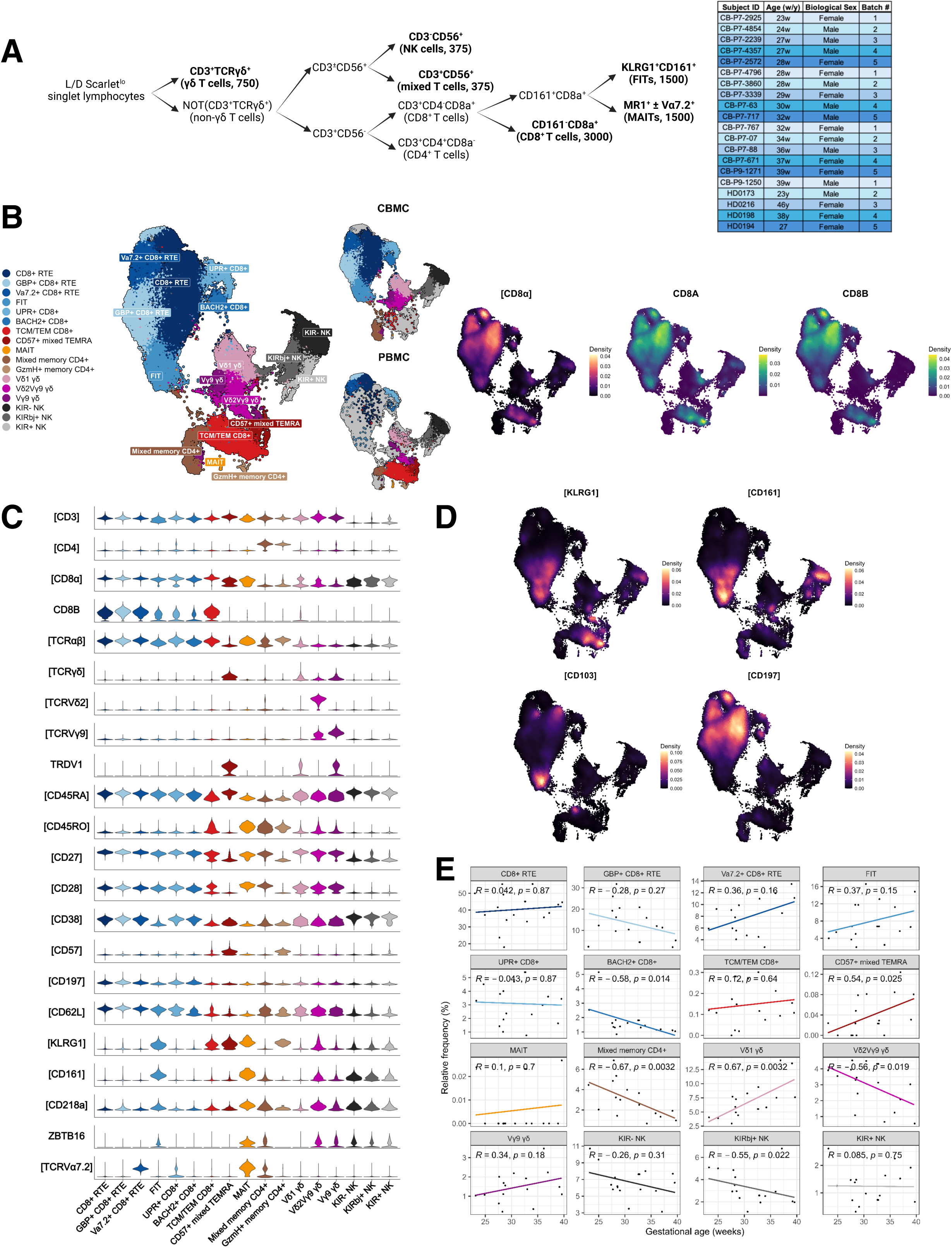
Infant naive CD8+ αβ T cells express innate-like features shared with other cord and peripheral blood lymphocytes. A) Sorting hierarchy employed for targeted population enrichment. Inset table shows subject age (w: weeks, y: years), sex and batch assignment. B) WNN UMAP with annotated cell clusters based on feature expression patterns as shown in Fig. 3C. Clusters are grouped by color according to their higher-order cell type i.e. naïve CD8α^+^ αβ T (blue), memory CD8α^+^ αβ T (red/orange), memory CD4^+^ αβ T (brown), γδ T (purple), or NK cells (grey) with subplots showing the distribution of CBMC and PBMC sample subsets. Inset density plots show the relative expression of CD8α protein or CD8A/B RNA. Feature names are stylized as ‘[PROTEIN]’ or ‘GENE’ to reflect their assay types. C) Stacked violin plots for each shown feature across cell clusters shown in Fig. 3B. All listed features correspond to background-normalized surface protein measurements except for the RNA features CD8B, TRDV1, and ZBTB16. D) Density plots for the listed surface protein features. E) Correlation plots relating cluster frequency to estimated gestational age across the infant cohort. Significance for linear models was determined via Spearman’s rank correlation test.

After excluding low quality cells, CD3^+^CD4^+^CD8^-^ cells and non-lymphocytes, we performed weighted nearest neighbor (WNN) integration of batch-corrected RNA and background-normalized protein datasets. Dimensional reduction and clustering yielded a UMAP graph segregated by cell type and memory phenotypes that recapitulated the expected enriched inputs with few batch-specific populations (Fig 3b, S3b, S3c). We identified 17 unique clusters among the merged neonatal and adult cells, which included both conventional CD8αβ^+^ T cells and non-conventional CD8α^+^ T cells (Fig 3b, 3c). Clusters were annotated based on surface antigen and marker transcript expression profile (Fig 3c).

Six clusters were identified as naïve conventional CD8^+^ T cells by their co-expression of CD8αβ, CD45RA, CD62L, CD27 and CD28 with low CD45RO expression (Fig 3c). As expected, naïve CD8^+^ T cell clusters were overrepresented by neonatal cells. Adult naïve CD8+ αβ T cells were present in all naïve clusters but were concentrated in two clusters (‘Naïve CD8+ RTE’, ‘Va7.2+ naïve CD8+ RTE’). Four clusters that corresponded to memory and effector CD8α^+^ T cells were predominantly derived from adults. Three γδ T and three NK cell clusters were detected in expected proportions based on the enrichment strategy. Vα7.2^+^ events were predominantly distributed across two clusters: a population of RTEs (CD38^+^CD45RA^+^) and a small but well-defined cluster of MAITs (CD8β^-^CD45RO^+^Vα7.2^+^MR1:5-OP-RU^+^) (Fig S3d). There was a small CD4^+^CD8a^-^CD45RO^+^ population (‘Mixed memory CD4+’) that was more frequent in the neonates and variably expressed ZBTB16 transcript and TCRγδ surface protein (Fig 3c).

Among the naïve T cell clusters, the putative FIT population (CD8αβ^+^CD45RA^+^KLRG1^+^CD161^+^) localized in UMAP space between CD38^+^ RTEs, CD8α^+^ effector/memory T cells and unconventional T cell clusters (Fig 3b, 3d). Compared to other naïve cells, FITs had lower CD197 (CCR7) expression, suggesting a reduced lymph node homing potential (Fig S3e). FITs also expressed more surface CD103 (ITGAE) than other naïve or memory CD8α^+^ T cells, recapitulating the cellular phenotype observed by flow cytometry (Fig 3d).

We next examined the relationship between cluster relative frequency and infant donor gestational age, in order to track the changing composition of cord blood lymphocytes during the second and third trimesters. Two γδ T cell clusters were positively (‘Vδ1’: R = 0.67, *p* = 0.0032) and negatively (‘Vδ2Vγ9’: R = -0.56, *p* = 0.019) correlated with gestational age (Fig 3e). The FIT cluster, while enriched in cord blood samples, was not significantly correlated with gestational age (R = 0.4, *p* = 0.11). The single MAIT cluster was absent from all but 3 infant donors and its frequency was not associated with age (Fig 3e).

The naïve fetal innate-like CD8αβ^+^ T cell transcriptome bridges conventional naïve and non-Conventional CD8a^+^ Lymphoid Cells.

To refine our FIT population definition and their predicted functional program, we derived per-cluster transcriptional markers by comparing each population to the rest of the sample to identify differentially expressed genes (Fig 4a). FITs expressed a set of transcripts distinct from other naïve CD8^+^ αβ T cells, which partially overlapped with transcripts expressed in γδ T cells (Fig 4a, 4b). KLRB1 (CD161) was upregulated in FITs, MAITs, Vδ2Vγ9 γδ T cells, and NK cells, reflecting its reported breadth of expression.^36–38^ FITs were enriched for multiple transcripts for surface receptors (i.e. NCR3, GPR183, DPP4) as well as the transcription factor MAF and the non-receptor tyrosine kinase BLK. NCR3 (CD337/NKp30), DPP4 (CD26), and MAF have been reported to be expressed by MAITs and Tc17 cells, but we detected only low NCR3 expression among the MAIT cluster relative to FITs.^39–41^ Additionally, FITs expressed a pattern of transcription factors that distinguished them from other naïve clusters (Fig 4b).

**Figure 4:**
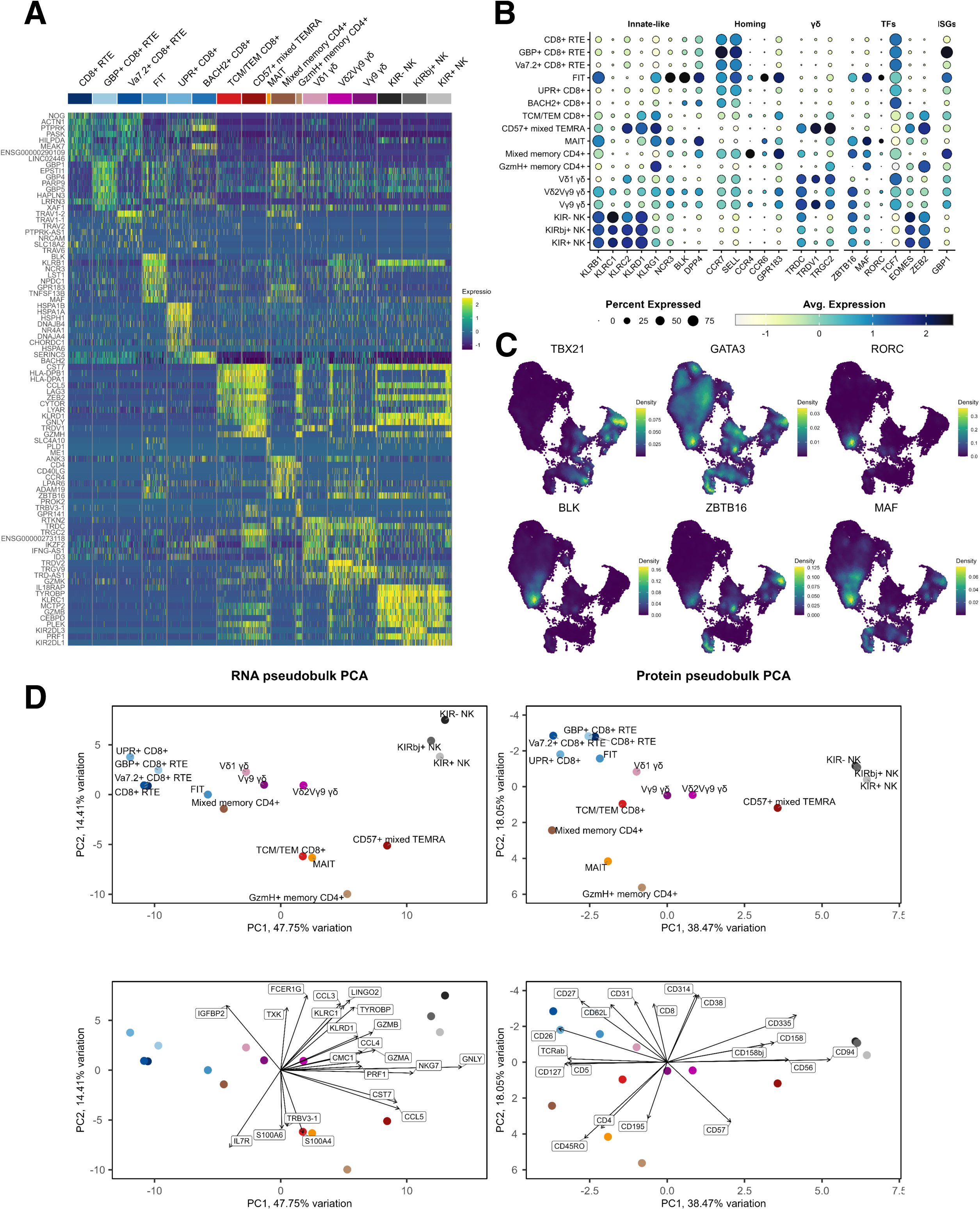
FITs express a unique set of transcriptional markers relative to other naïve CD8α^+^ αβ T cells that are shared by other innate-like T cells. A) Heatmap showing diqerential expression of transcriptional markers for each cluster versus all other cells. Expression levels are scaled by z-score for clarity. B) Dot plot contrasting expression patterns for listed transcriptional markers (columns) for each cell cluster (rows). Per-cluster median scaled expression levels are shown for each feature. C) Feature expression shown via density plots for selected transcription factors. D) Principal components analysis plots showing the relative positioning of cluster RNA and protein pseudobulk profiles. Bottom plots for both assay types show the PC1 and PC2 loadings for the top 15 variable features.

We further examined the expression pattern of key T helper cell transcription factors (i.e. TBX21, GATA3, RORC) as well as BLK, ZBTB16 (PLZF), and MAF.^42,43^ (Fig 4c). Like all other naïve CD8α^+^ αβ T cell clusters, FITs lacked expression of TBX21 (T-bet) and were positive for GATA3. However, FITs were unique among naïve CD8α^+^ αβ T cells in their expression of RORC, BLK, MAF and ZBTB16. BLK is associated with B cell development and differentiation but has also been shown to be expressed in human thymocytes and is required for the development of murine IL-17-producing γδ T cell populations.^44,45^ Similarly, c-MAF is broadly expressed by T cell subtypes but is also notable for its STAT3-dependent role in inducing RORγt expression among human and murine Th17 and IL-17^+^ γδ T cells.^46–48^

CD8α^+^ subpopulations can be located along an adaptive to innate spectrum, and we hypothesized, based on their innate-like surface phenotype, that FITs would share a transcriptome with other innate CD3^+^ and CD3^-^ lymphoid populations.^49^ To address this hypothesis, we performed separate pseudobulking with RNA and non-hashing protein features to derive an aggregate expression profile for each cell cluster. Principal components analysis of both data types yielded a distribution of clusters that reflected cell type annotations (Fig 4d). The grouping of cell types was generally similar between the RNA- and protein-specific PC plots, indicating that the surface phenotypic markers used for discriminating populations generally reflected differences in cells’ gene expression. PC1 of the RNA-based UMAP stratified cells along an adaptive-to-innate axis, as reflected by positive PC1 loadings for NK-enriched and cytotoxicity-related transcripts (e.g. GNLY, PRF1) and negative PC1 loadings related to T cell proliferation and survival (i.e. IL7R, IGFBP2) (Fig 4d). FITs were adjacent to other naïve CD8^+^ T cell clusters in both PC plots but located closer to innate lymphoid populations along PC1, consistent with their unique transcriptome relative to other naïve clusters.

### Fetal Innate-like T cells Exhibit a DiFerential Transcriptional Profile Relative to Conventional CD8α^+^ ab T cells and are Clonally Diverse

Murine neonatal T cells have been shown to have a greater capacity for effector, rather than memory, differentiation compared to their adult counterparts.^20,50^ Consistent with these findings, the KLRG1^+^CD161^+^ population (‘FITs’) we found in cord blood shared multiple phenotypic and transcriptional markers with the small population of MR1-reactive TCRVα7.2^+^ MAITs, which are known to be highly plastic and often described as possessing an “effector memory” phenotype in circulation.^51,52^ We thus hypothesized that the putative FIT cluster may harbor a transcriptional signature of an atypical naïve CD8α^+^ T cell differentiation state. However, prior studies using CD8α as a broad marker of conventional CD8^+^ T cells without combinatorial measurement of CD8β, RNA, protein features and TCR sequences were unable to reliably align fetal CD8α^+^ T cells to either conventional or unconventional T cell subsets, and may even have misidentified what was truly an innate lymphoid population that has upregulated CD8α.^53^

To rigorously delineate the multiple groups of conventional αβ CD8^+^ T cells, we restricted our analyses to those cells with a surface phenotype annotated as "naïve CD8^+^" (CD4^-^CD8ab^+^CD45RA^+^CD45RO^-^CD27^+^CD28^+^) or "memory CD8^+^" (CD4^-^CD8ab^+^CD45RA^+/-^ CD45RO^+^) (Fig S3b). We further excluded cells that expressed TCRγδ surface protein or transcript or that lacked surface expression of TCRαβ. Reintegration with RNA and protein features yielded cell clusters that could be identified by expression of typical surface protein markers across both assay types and were segregated in UMAP space by naïve and memory/effector status (Fig 5a, Fig S4a). Naïve CD8 T cell clusters were detected among cord blood (RTEs and FITs) or adult donors (GBP+ adult naïve and adult naïve), while all effector and memory clusters were overrepresented among adult donors (Fig 5a). Previously identified MAITs were distinguished from the closely associated GZMK+ FIT cluster by transferring prior annotations of this small number of cells. Notably, the GZMK+ FIT cluster expressed transcriptional features frequently associated with MAITs (e.g. SLC4A10, CEBPD, GZMK) but lacked MR1 tetramer reactivity and did not express the Vα7.2 TCR side chain (Fig S4b).

**Figure 5:**
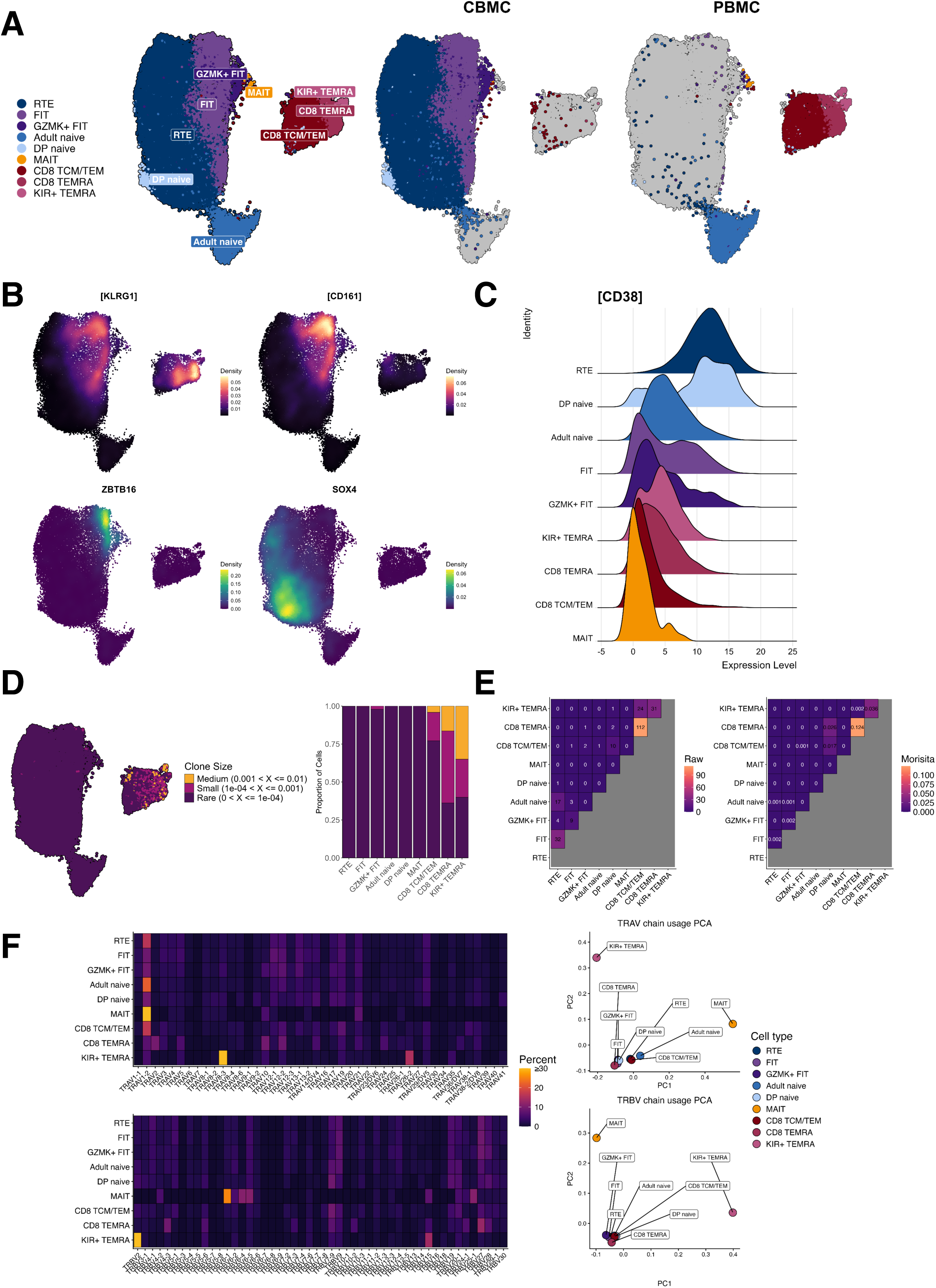
FITs are a unique and clonally diverse subpopulation among conventional infant and adult CD8α^+^ αβ T cells. A) WNN UMAP generated from clusters previously annotated as ‘CD8+ naïve’ or ‘CD8+ memory’ after excluding any events with positive surface expression of TCRγδ or CD4. B) Feature expression shown via density plots for selected protein or RNA features. Feature names are stylized as ‘[PROTEIN]’ or ‘GENE’ to reflect their assay types. C) Ridge plots contrasting CD38 surface protein expression density among the annotated cell populations. D) Clone size estimates for TCRα/TCRβ CDR3 amino acid clonotypes identified in each annotated cell population. Stacked barcharts show the relative proportion of cells with diqerent clone size estimates among each population. E) Quantification of clonal overlap of TCRα/TCRβ CDR3 nucleotide clonotypes between specified CD8^+^ αβ T cell populations. The inset number shown for pairwise comparisons between clusters corresponds to raw number of shared clones (left) or the Morisita-Horn index (right) for sequenced clones found to be shared between clusters. F) Summary heatmaps showing relative frequency of TRAV (top) and TRBV (bottom) chains among annotated cell populations. Associated principal components analysis plots show the relative arrangement of cell populations based on their TRAV (top) and TRBV (bottom) chain usage pattern.

Surface expression of KLRG1 and CD161 was observed on both naïve (’FITs’) and memory CD8^+^ (‘CD8 TCM/TEM’) clusters with a subset of FITs expressing the transcription factor ZBTB16, as previously shown (Fig 5b). Conversely, FITs expressed lower levels of SOX4, a transcription factor enriched among RTEs and known to promote memory differentiation, compared to other infant naïve CD8α^+^ T cells (Fig 5b).^42^ Together, these results suggest that FITs may have a unique differentiation program, including reduced memory potential, compared with other naïve populations. Surface CD38, a marker of recent thymic emigration, showed a gradient from maximal in RTE CD8α^+^ T cells to only more sparsely expressed in FITs (Fig 5c).^42^ Thus, CD38^lo/int^ FITs either emerge from the thymus with lower CD38 expression or downregulate its expression following a longer time spent in peripheral circulation.

We next examined whether FITs harbor a clonally expanded TCR repertoire, as is seen in memory or effector populations, by performing per-cell type clone size estimates. Memory and effector CD8α^+^ T cell showed an expected signature of clonal expansion (Fig 5d). Like other cord blood RTEs and adult naïve cells, FITs contained exclusively small or rare (<0.1%) clonotypes (Fig 5d), indicating an expected polyclonal population for naïve cells. Additionally, comparison of TRA/TRB clonotypes between FITs and other cell clusters revealed minimal overlap in TCR repertoires except among memory CD8α^+^ T cells (Fig 5e). We further examined TRAV and TRBV usage frequencies to test whether FITs exhibited a semi-invariant TCR repertoire reminiscent of MAITs (Fig 5f). While Va7.2^+^ events were overrepresented in this dataset as a result of our sort strategy, we found that all naïve CD8α^+^ T cells expressed a broad range of TRAV and TRBV chains, in contrast to the more restricted chain usage seen among MAIT and KIR+ TEMRA cells. Finally, we did not observe any differences in CDR3 length and 3-mer nucleotide usage frequency between naïve CD8α^+^ T cell populations or any donor-specific TRAV/TRBV chain usage frequencies among FITs (Fig S4c-e). Collectively, these results demonstrate that FITs are neither clonally expanded from other naïve populations nor are they likely to be derived from the semi-invariant and public clones commonly seen in other innate lymphoid populations such as MAITs. Further, FITs’ TCR repertoire was did not differ in CDR3 length or sequence composition relative to infant RTEs, suggesting that their phenotype does not directly result from an alternative process of TCR-specific thymic selection, as has been shown for MAITs.^54,55^

We next performed gene ontology (GO) gene set enrichment analysis (GSEA) to determine differential gene pathway expression between FITs and other conventional naïve CD8ab T cells. Relative to infant RTEs, FITs were enriched for GO terms associated with adaptive immune responses, cytokine signaling, and inflammation (Fig 6a, Fig S5a). Relative to adult naïve cells, FITs were enriched for terms associated with adaptive immune responses, cell killing, and chemotaxis (Fig 6b, Fig S5b). Despite these transcriptional indicators of an effector response, FITs expressed few transcripts related to cytokines or cytotoxicity except for the GMZK-expressing cluster (Fig 6b). Instead, they could be distinguished from infant RTE and adult conventional naïve CD8^+^ T cell populations by their expression of diverse transcripts related to innate-type functions (e.g. NCR3, CR1), type 17 immunity (e.g. RORC, IL23R) and mucosal homing potential (e.g. CCR4, CCR6, CXCR6). Additionally, we performed transcription factor activity inference analysis to determine whether cellular clusters were enriched for specific transcription factors and their downstream targets.^56^ FITs upregulated a set of regulons shared with adult memory CD8α^+^ T cells but not infant RTEs or adult naïve cells (e.g. JUN, FOSL1, STAT4) and lacked expression of the TCF7 regulon, unlike other naïve cells (Fig 6c).

**Figure 6:**
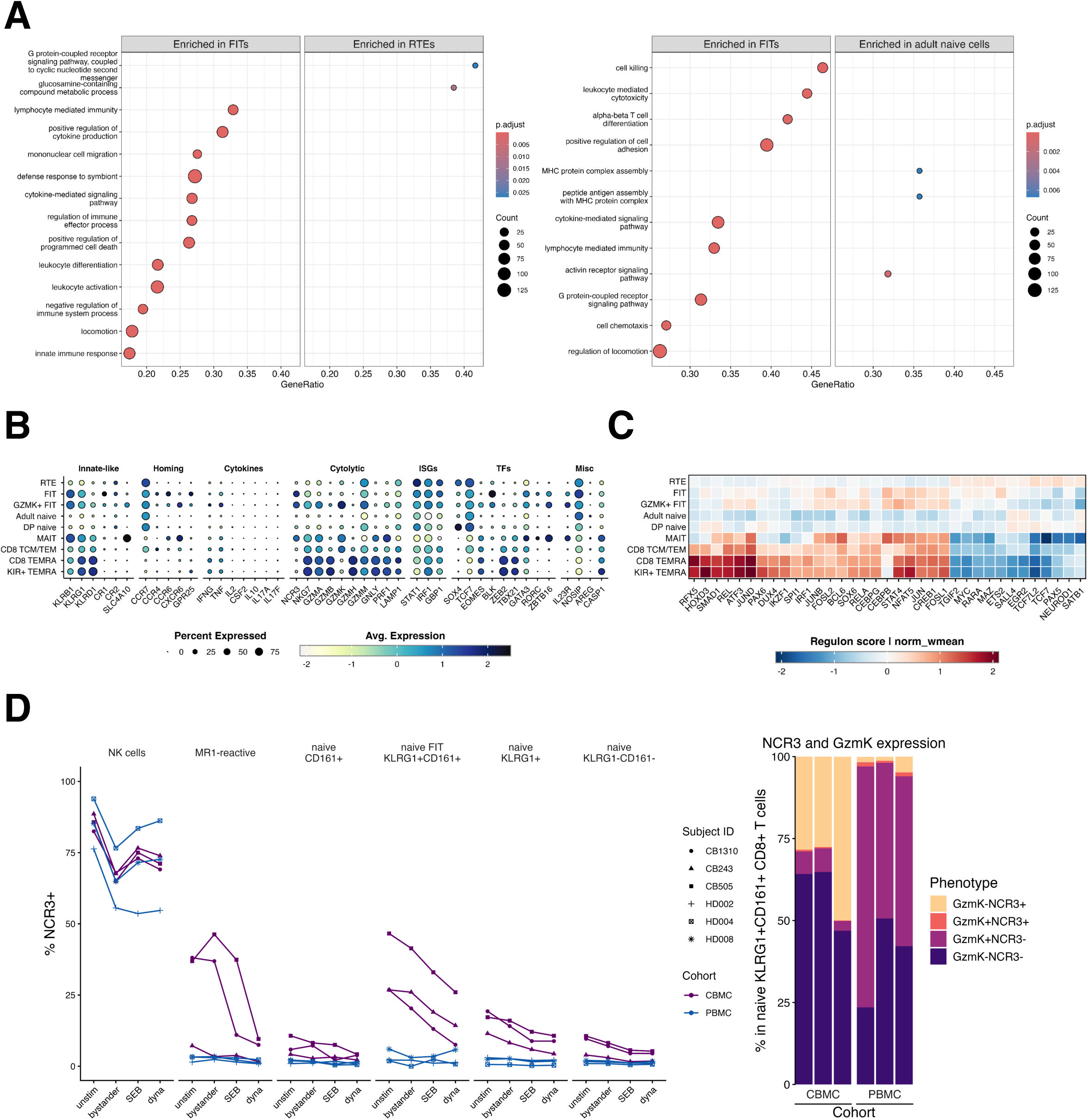
FITs are enriched for transcripts related to immune eWector processes but lack expression of canonical cytokine or cytolytic transcripts. A) GO GSEA significance plots derived from comparisons between FIT and infant RTE (left) and FIT and adult (right) naïve CD8α^+^ αβ T cell clusters. Shown GO terms represent a curated list of significant terms after removing redundant entries. B) Dot plot contrasting expression patterns for listed transcriptional markers (columns) for each conventional αβ CD8+ T cell cluster (rows). Per-cluster median scaled expression levels are shown for each feature. C) Heatmap summarizing estimated regulon scoring among conventional CD8+ αβ T cell clusters (rows) for the specified transcription factors (columns). Transcription factor activity inference was performed using decoupleR and dorothea. D) Quantification of NCR3 surface expression among full-term infant (n=3) and adult (n=3) lymphocyte populations under specified stimulation conditions. Stacked barcharts show the expression patterns of NCR3 and GzmK among naïve KLRG1^+^CD161^+^ CD8α^+^ T cells for individual subjects. ‘unstim’ = media control, ‘bystander’ = IL-12/IL-18, ‘SEB’ = ‘Staphylococcal Enterotoxin type B’, ‘dyna’ = ‘anti-CD3/anti-CD28 beads’

Considering NCR3 and GZMK as markers that could further refine our FIT phenotype, we used flow cytometry to compare NCR3 (NKp30) and granzyme K protein expression in stimulated and unstimulated conditions; CBMC and PBMC samples were phenotyped following bystander cytokine (i.e. IL-12/IL-18) and TCR-specific stimulation (i.e. SEB, anti-CD3/CD28 Dynabeads). Relative to infant KLRG1^-^ and all adult naïve CD8α^+^ T cells, baseline NCR3 expression was greater in KLRG1^+^ naïve infant T cells, and highest in FITs (Fig 6d, Fig S5b). Cytokine stimulation resulted in downregulation of NCR3 among CD8α^+^ NK cells and conventional naïve CD8α^+^ populations, but was less consistently observed among MR1-reactive T cells. NCR3 expression was decreased following TCR-dependent stimulation in all populations except NKs (Fig 6d). Further, NCR3 and granzyme K were rarely co-expressed on naïve KLRG1^+^CD161^+^ CD8α^+^ cells, supporting the functional heterogeneity we observed via single cell transcriptomic profiling (Fig 6d). These results suggest that NCR3 is expressed by neonatal naïve CD8a^+^ T cells and is highest in KLRG1^+^ populations, especially in resting FITs.

### Functional and Phenotypic Profiles of Fetal Innate-like T Cells Relative to Recent Thymic Emigrants and Adult Naïve CD8α^+^ T cells Following Stimulation

Prior evidence from mice and humans led us to hypothesize that fetal innate-like CD8α^+^ αβ T cells should be competent to respond to bystander cytokines IL-12 and IL-18.^22,49^ However, intracellular cytokine staining of IL-12/IL-18-stimulated KLRG1^+^CD161^+^ CD8^+^ αβ neonatal T cells resulted in minimal detectable IFNγ, TNF, or IL-8 by cytometry, regardless of gestational age at birth. To determine if there was any measurable response by FITs to bystander stimulation based on gene expression, we evaluated which clusters differed in their expression of cytokine transcripts between stimulated and unstimulated samples. Separate visualization by stimulation condition revealed an expected upregulation of IFNγ transcript in CD8αα^+/-^ NK, Vδ1 and Vδ2Vγ9 γδ T clusters (Fig S6a). Very few naïve CD8α^+^ αβ T cells expressed IFNG, CXCL8 or IL17A transcripts in response to cytokine stimulation. However, FITs did show slight upregulation of TNF, consistent with flow cytometry results after PMA/ionomycin stimulation (Fig 7a), raising the potential for either direct activation from PMA/ionomycin or indirect activation through other innate-derived signals in that condition.

**Figure 7:**
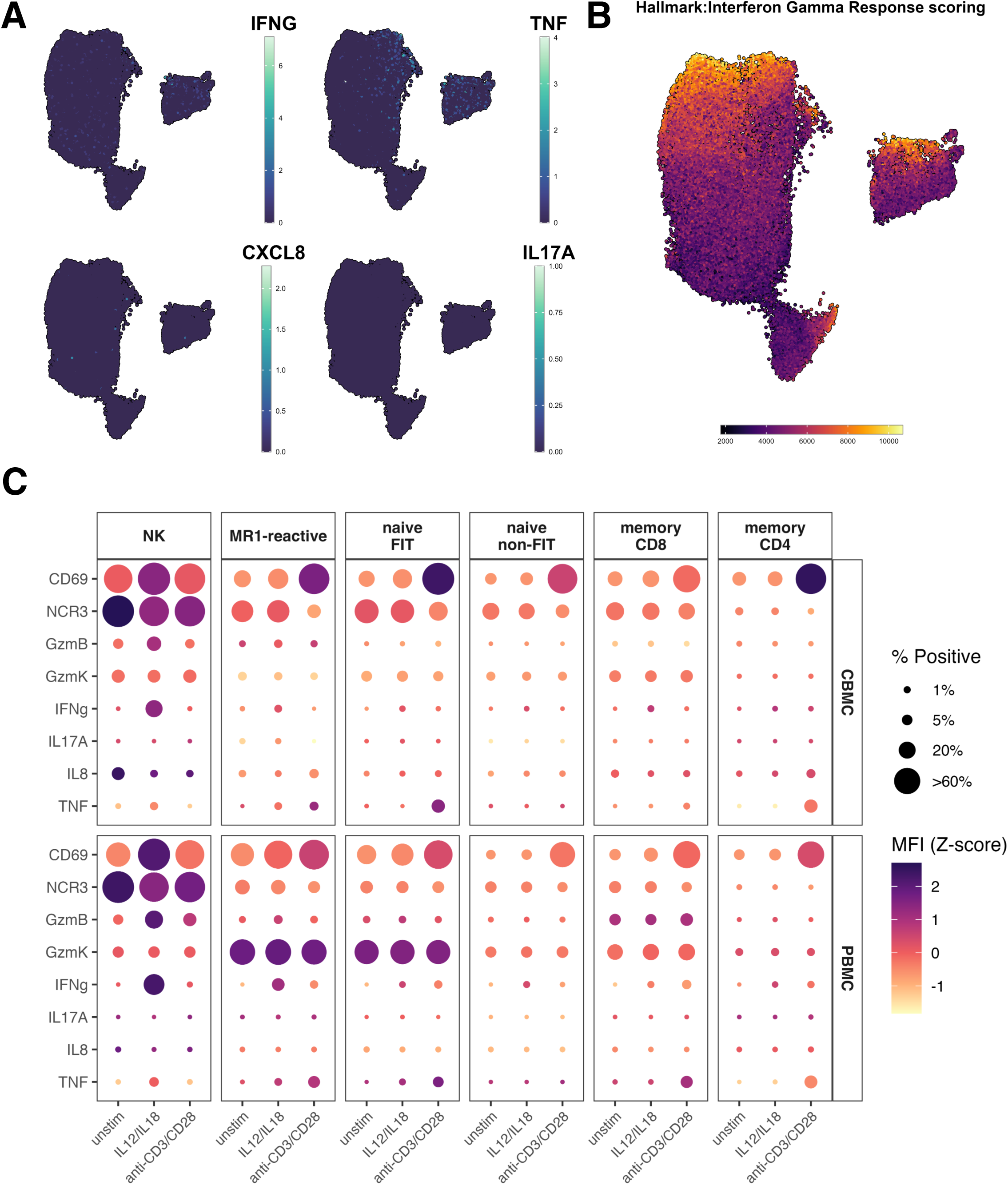
Infant CD8α^+^ αβ T cells appear minimally capable of bystander activation but express an IFNγ-response gene set. A) Feature plots showing detection of listed eqector transcripts among conventional CD8α^+^ αβ T cells. Shown UMAP was derived from both unstimulated (i.e. media alone) and bystander-activated (i.e. IL-12/IL-18-stimulated) samples. B) Visualization of single cell GSEA scoring results for conventional CD8^+^ αβ T cells using the ‘Hallmark Interferon Gamma Response’ gene set. Scoring was performed using the escape package. C) Dot plots showing % positivity (relative to parent population) and median fluorescent intensity for surface and intracellular eqector molecules under shown stimulation conditions with n = 3 subjects per group. MFI values are z-scaled per row for ease of visualization. Phenotypic populations were manually gated as follows: NK cells – CD3^-^CD56^+^, MR1-reactive – CD3^+^CD8α^+^MR1:5-OP-RU^+^, naïve FIT – CD3^+^CD8α^+^CD45RA^+^CD27^+^CD161^+^KLRG1^+^, naïve non-FIT - CD3^+^CD8α^+^CD45RA^+^CD27^+^CD161⊕KLRG1, memory CD8 - CD3^+^CD8α^+^CD45RA⊕CD27, memory CD4 - CD3^+^CD4^+^CD45RA⊕CD27.

Multiple infant and adult naïve CD8^+^ αβ T cells clusters expressed GBP1, an IFNγ-inducible GTPase important for controlling intracellular infections as well as diverse cell-intrinsic immune responses (Fig 6b).^57^ To assess if naïve and FIT activation could be downstream of IFNγ secreted by classically innate populations, we scored individual cells using the Hallmark Interferon Gamma Response gene set and visualized enrichment scores by UMAP. Both naïve and memory CD8^+^ αβ T cells appeared to upregulate IFNγ responsiveness genes under bystander activation conditions even if this did not result in cytokine transcript expression (Fig 7b). Additionally, GBP1+ FITs were enriched for GO terms associated with cytokine signaling and antiviral innate immune responses (Fig S6b), consistent with an interferon-responsive program.

We next measured expression of key cytokines and cytolytic molecules following IL-12/IL-18 and anti-CD3/CD28 stimulation of CBMC and PBMC lymphocytes to confirm these findings via flow cytometry (Fig 7c). NK cells upregulated CD69 and expressed IFNγ under bystander activation conditions, as did adult but not infant MR1-reactive CD8α^+^ T cells. A subpopulation of infant and adult naïve KLRG1^+^CD161^+^ CD8α^+^ T cells responded to TCR-dependent stimulation by producing TNF, but none of the other measured cytokines (i.e. IFNγ, IL-17A, IL-8) were detected among these cells. Expression of cytolytic molecules was heterogeneous among CD8α^+^ lymphocytes in that granzyme B was most highly expressed by NK cells of all ages and adult memory CD8α^+^ T cells, while granzyme K was detected in a subset of infant KLRG1^+^CD161^+^ CD8α^+^ T cells as well as adult MR1-reactive and memory CD8α^+^ T cells.

In total, these results suggest that previously observed infant naïve CD8α^+^ T cells’ responsiveness to bystander activation conditions may be due to secondary IFNγ or other indirect signaling rather than via a direct and cell-intrinsic IL-12/IL-18 signaling cascade. This may be particularly true under stimulation conditions involving co-culture with mononuclear cells that share similar surface profiles (e.g. CD8a^+^ MAITs, NK and γδ T cells). Further, despite FITs’ expression of transcription factors related to type 17 differentiation, they do not appear poised at the time of birth to produce type 17 cytokines under the stimulation conditions tested here.

## Discussion

The previously held paradigm that human neonatal adaptive immunity is maintained in a suppressed state has more recently been challenged by several publications demonstrating a unique gene regulatory and functional landscape of CD8α^+^ T cell populations in early life.^21,22^ These findings have been robust to different *in vitro* and *in vivo* approaches, but previous studies have not addressed the influence that known CD8α-expressing T and innate lymphoid cell populations could have on these results. Putative fetally-derived subsets therefore require a refined definition that considers phenotypic and functional overlap with other CD8α-expressing populations. Indeed, recently published guidelines for T cell nomenclature in an age of high-dimensional cellular phenotyping have emphasized the need for terminological specificity when describing previously monolithic populations (e.g. ‘CD8^+^ T cells’).^58^ By using integrative, multi-omic profiling of CD8aα- and CD8ab-expressing lymphocytes, we provide evidence that the CD8α^+^ T cell compartment in neonates is highly heterogeneous and includes both conventional and unconventional T and NK cell populations. We further show that cord blood is enriched for a population of naïve KLRG1^+^CD161^+^ CD8αβ^+^ T cells (‘FITs’), which exhibit a unique transcriptomic and phenotypic profile that shares some but not all features with well-described innate-like populations.

While FITs can be readily distinguished from γδ T and NK cells, they bear a close resemblance to MAITs in their surface phenotype and expression of key transcription factor genes. By using tetramer reagents to identify MR1-reactive and/or semi-invariant T cells (i.e. CD8a^+^ TCRVα7.2^+/-^MR1:5-OP-RU^+^), we found that in contrast with MAITs, cord blood-derived KLRG1^+^CD161^+^ FITs expressed the CD8αβ heterodimer, were not MR1 tetramer reactive, and were distinguished by their co-expression of naïve phenotypic markers (i.e. CD45RA^+^CD27^+^) and NCR3 (NKp30). Furthermore, FITs are clonally diverse and lack the invariant TCRa chain enrichment characteristic of innate lymphoid populations, like their non-FIT naïve CD8α^+^ T cell counterparts. MAITs were exceedingly rare in circulation in both preterm and full-term infants, consistent with the reported postnatal expansion and persistence of MAITs in childhood and throughout the adult lifespan.^17,54,55,59^ CD161 itself, which is often used as a surrogate marker of MAITs in adult circulation, was expressed by multiple T cell populations in early life, including MAITs, γδ T and NK cell populations. Previous studies have shown that CD161^+^ MAIT and non-MR1-reactive CD8αα^+^ as well as IL-17^+^ CD8α^+^ T cells emerge from a pool of polyclonal CD8αβ^+^ αβ T cells marked by CD161 expression, making it even more important to identify antigen combinations that more reliably isolate the fetal CD8a^+^ T cell signature.^59–61^

The FIT population could be detected in fetal circulation beginning at 23 weeks of gestation, indicating that FITs arise *in utero* during or prior to the late second trimester. By leveraging recent studies of CD38 as a specific marker for CD8α^+^ T cell RTEs, we can speculate that the lower expression of CD38 on KLRG1^+^CD161^+^ CD8α^+^ T cells reflects their earlier thymic egress and maturation during circulation.^42^ It is presently unclear whether FITs emerge from the thymus with an innate-like transcriptome and phenotype or whether they acquire these features after a longer period of peripheral circulation. Relative to their naïve RTE counterparts, FITs exhibited higher expression of ZBTB16(PLZF), MAF and BLK, but downregulated transcription factors important for maintenance of the naïve cell state or true memory formation (e.g. TCF-1, SOX4, respectively). This finding is counter to studies using adult animal models that show epigenetically-controlled suppression of PLZF in non-innate T cell populations.^62^ A pair of recent reports have shown that fetal and infant mucosal tissue resident αβ T cells, while predominantly naïve, express a transcriptome similar to FITs, including PLZF-regulated transcripts.^63,64^ FITs may therefore possess a gene regulatory landscape closer to so-called “preset T cell” populations than to conventional naïve CD8α^+^ αβ T cells, enabling them to undergo a conventional differentiation pathway while executing an innate-like effector program.^8,63–65^ Interestingly, FITs were unique among naïve CD8αb^+^ T cells in their enhanced expression of NCR3 transcript and surface protein. NCR3 is an NK cell receptor with diverse ligands and with known roles in recognition of tumor or infected cells, cytotoxicity and human cytomegalovirus immune evasion.^66,67^ NCR3 is induced in umbilical cord blood CD8α^+^ T cells following IL-15 signaling and enhances cytokine secretion and cell killing *in vitro*.^68^ Whether NCR3 interacts with its diverse endogenous ligands *in utero* and how this might influence perinatal T cell functions *in vivo,* including tumor surveillance during dynamic fetal development, remains to be established.

Despite increased resting expression of CD218a (IL18R1), FITs were minimally capable of bystander activation, as indicated by the absence of IFNG transcript expression or intracellular cytokine detected among these cells following IL-12/IL-18 stimulation.

These results highlight the importance of deploying a refined phenotype to distinguish subpopulations among heterogeneous CD8α-expressing cells, particularly when performing *in vitro* manipulation of bulk or mixed lymphocyte populations. We identified subsets of FITs and infant RTEs as well as adult naive and memory CD8α^+^ T cells that appeared to respond to neighboring cell-derived IFNγ. It is likely that upregulation of specific members of the broad class of interferon-stimulated genes reflects secondary as opposed to direct IL-12 and IL-18 reactivity. Intriguingly, interferon-induced GBP family members, especially FIT-enriched GBP1, are receiving renewed attention as mediators of cell-autonomous immunity against intracellular pathogens. GBP1’s role in inflammasome assembly and pyroptosis, as well as its capacity for forming a multimeric "capsule" on the Gram-negative bacterial cell surface via direct binding to LPS, points to another potential fetal CD8ab^+^ T cell innate immune function.^69,70^ Several groups have recently reported the enhanced expression of GBP transcripts among fetal compared to postnatal infant intestinal T cells and in inflammatory conditions such as intrauterine infection.^63,71^

FITs produced robust amounts of TNF cytokine following PMA/ionomycin and TCR-specific stimulation, which typically entails a pro-inflammatory response in the postnatal context. However, T cell-derived TNF is also known to be important for normal fetal intestinal development.^72^ Whether FITs can be recruited to homeostatic or inflamed mucosal sites is unknown, as is their site-specific effector profile, but their activation and TNF secretion during fetal development or postnatal life could play either a protective or pathologic role. We detected minimal expression of transcripts encoding other potential CD8α^+^ αβ T cell proinflammatory cytokines (e.g. IFNγ, IL-8, IL-17) under our experimental conditions. Although FITs were enriched for transcripts associated with type 17 immunity (e.g. RORC, CD161, IL23R), we did not detect upregulation of IL17A/F transcripts or production of IL17A cytokine following T cell stimulation. These findings are in agreement with older studies reporting that KLRG1 is expressed abundantly among cord blood naïve CD8α^+^ T cells and marks a population with enhanced proliferative capacity but that lacks the ability to secrete IFNγ, perhaps reflecting KLRG1’s capacity to inhibit TCR signaling when bound to epithelial E-cadherin.^73,74^ Notably, KLRG1 is used typically as an experimental marker of T cell effector differentiation and/or exhaustion and is rarely expressed on circulating conventional naïve T cells outside of the perinatal period.^75–77^ Finally, FIT subpopulations expressed few or undetectable levels of cytotoxic/cytolytic transcripts except for the GZMK^+^ FIT subcluster, which co-clustered with bona fide adult circulating GZMK^+^ MAITs but lacked essential features of this well-described innate-like population (i.e. TCRVα7.2 expression or MR1:5-OP-RU tetramer reactivity). Even in the absence of a clear cytotoxic transcriptional profile, Granzyme K may instead initiate the complement cascade, oCering yet another mechanism in neonates to initiate and propagate an innate immune response under otherwise tolerogenic conditions.^78,79^

### Limitations

Our study has specific limitations that should be considered. While we were able to use cord blood from infants of varying gestational age to investigate peripheral T cells across the third trimester, paired postnatal infant PBMC samples would clarify whether fetal innate-like CD8αβ^+^ αβ T cell populations persist in circulation after birth. Similarly, unperturbed fetal or longitudinal postnatal infant samples will be helpful to explore the apparent FIT program and its link to processes initiated by prematurity and/or labor. Growing evidence suggests that perinatal immune functions are highly tissue-specific and distinct from typical adult mucosal immune responses.^63,64,80–82^ As such, it will be important for the field to conduct comparative studies of both infant blood and tissue to clarify whether circulating FITs bear resemblance to those found at barrier sites, which are most likely to be involved in acute pathogen control, and which participate in a recall response. In addition, if FITs are found to be present beyond infancy, tumor models will be useful to explore their potential role in anti-tumor activity given their co-expression of several antigens known to associate with anti-tumor effects.^66,83^

### Conclusion

The developing fetal immune system balances the competing demands of organogenesis, immune tolerance, and pathogen control while assembling diverse and unique layers of cellular protection that will ease the transition to postnatal life. Here, we report that infant cord blood contains naïve CD161^+^KLRG1^+^ CD8αβ^+^ αβ T cells, which can be distinguished from other naïve CD8α^+^ T cell populations by surface phenotype alone. Relative to other naïve CD8α^+^ T cells, FITs express diverse transcriptional features related to mucosal homing and effector functions but lack overt expression of classical cytotoxicity-related transcripts. FITs can be further subdivided by their expression of diverse transcripts known to be regulated by PLZF as well as a set of interferon-stimulated genes. While FIT clusters bear phenotypic resemblance to well-characterized UTC lineages, they are polyclonal and thus not simply precursors for these cell types. In total, their unique transcriptomic features and effector profile further reinforce the potential for FITs to be poised to serve a broad array of functions from pathogen responsiveness, to organogenesis, to tumor surveillance in the singularly dynamic period of fetal development and birth. Future studies will be aimed at determining when, where and how FITs emerge *in utero* while exploring their postnatal distribution, differentiation and functionality.

## Methods

### Clinical Sample Procurement

Umbilical cord blood samples were collected according to University of Rochester Medical Center (URMC) Institutional Review Board protocols (Study #00000729) and processed according to previously published protocols before cryopreservation. Healthy donor adult peripheral blood samples were collected according to URMC IRB protocols (Study #00000009). Briefly, cord blood (CBMC) and peripheral blood mononuclear cells (PBMC) were isolated via Ficoll/Percoll density gradient preservation before enumeration and cryopreservation.

For flow cytometric assays, de-identified cryopreserved CBMC samples were obtained from the URMC Umbilical Cord Blood Repository. CBMC samples were selected according to the following criteria: no clinical or histologic evidence of maternal chorioamnionitis, pre-term prelabor rupture of membranes (PPROM) or pre-eclampsia; and singlet infant subjects without known congenital abnormalities. PBMC samples were selected according to the following criteria: adult subjects between 18 and 60 years of age with no known chronic conditions or reported infections in the prior 2 weeks.

For single cell sequencing assays, cryopreserved CBMC were selected according to the above criteria together with the following additional criteria: no antenatal maternal steroid use; APGAR > 4 at 5 minutes; and sex matching by gestational age (GA) when possible. Available donor metadata are shown in Table S1.

### Sample Thawing & Stimulation

Cryopreserved vials were thawed in batches of ≤5 subjects in a 37°C water bath for 1-2 minutes until a small fragment of ice remained. Thawed cells were washed with pre-warmed R10 (RPMI-1640 with 10% FBS & 1X Pen/Strep; Corning, cat. #10-04-CV; Seradigm, cat. #1500-500; ThermoFisher, cat. #15140122) supplemented with 1X Universal Nuclease (Thermo Scientific, cat. #88702) (’thawing medium’) and combined in a 15 mL conical tube. Samples were centrifuged for 5 min at 300g and 25°C and the supernatant was decanted. Cell pellets were resuspended in 37°C thawing medium and re-centrifuged as above. CBMC samples were treated with PharmLyse (BD, cat. #555889) according to manufacturer recommendations to lyse contaminant erythrocytes before washing with PBS (pH 7.2, Gibco, cat. #20012027) and re-centrifugation as above. Cell pellets were resuspended in 2mL warm R10 medium before cell number and viability were measured using AOPI assay settings on the K2 Cellometer (revvity, cat. #CMT-K2-MX-150). Samples were rested overnight at 37°C and 5% CO2 before remeasurement of cell number and viability the following morning.

Only samples with ≥80% final cell viability were used for subsequent experimental assays. Viable samples were resuspended in warm R10 at 10e6 cells/mL before plating at 1e6 cells/well in a 96-well V-bottom plate (Corning, cat. #3-9684)

Plated cells were stimulated with either 1) PMA (500 ng/mL, Sigma, cat. #P8139) and ionomycin (1 μg/mL, Sigma, cat. #I0634) for 6 hours (’P/i-treated’), 2) IL-12 (p70) (10ng/mL, Biolegend: cat. #573008) and IL-18 (10ng/mL, Biolegend, cat. #592104) for 18 hours (’bystander activated’), 3) anti-CD3/anti-CD28 Dynabeads (1:2 bead:cell ratio, Gibco, cat. #11132D) for 10 hours (‘CD3/CD28 stimulated’), 4) Staphyloccocal Enterotoxin Type B (2μg/mL, List Labs: cat #122) for 10 hours (‘SEB stimulated’) or 5) R10 medium alone for 18 hours (’unstimulated’). GolgiStop (BD, cat. #554724) and GolgiPlug (BD, cat #555029) were added for the final 4 (‘P/i-treated’) or 8 hours (all other conditions) of each incubation interval to preserve a cell-specific cytokine profile for intracellular staining.

### Immunostaining: Flow Cytometry

Stimulated cells were centrifuged for 5 min at 300g and 25°C and the supernatant was discarded using a multichannel pipette (’live cell washing’). Cells were washed twice with PBS with 2% FBS (’staining buCer’), once with PBS, and stained with LIVE/DEAD Fixable Blue Dye (1:200 dilution in PBS with 0.5% FBS, Invitrogen, cat. #L23105) for 10 min at room temperature. Fc receptor blocking was performed with Human TruStain FcX (2.5 μL/well in PBS with 4% FBS, Biolegend, cat. #422305) for an additional 10 min at room temperature. PE-conjugated MR1:5-OP-RU tetramers (1:500 dilution in staining buCer, NIH Yerkes Core) were added to each well before incubation for 10 min at 4°C.

Cocktails of antibodies targeting surface antigens were prepared using previously established optimal reagent concentrations in Horizon Brilliant Staining BuCer (BD, cat. #563794) and added to each well (final staining volume: 100 μL) before incubation for 20 min at room temperature. Blocked and surface-stained samples were washed twice with staining buCer and were then fixed and permeabilized with the Cytofix/Cytoperm Fixation/Permeabilization kit (BD, cat. #554714) according to manufacturer recommendations and with subsequent centrifugations at 800g. Cocktails of antibodies targeting intracellular antigens were prepared as above using Perm/Wash BuCer as a solvent and added to each well (final staining volume: 50uL) before incubation for 30 min at room temperature. Fully stained cells were washed twice with Perm/Wash BuCer before resuspension in 150uL staining buCer for subsequent acquisition. A full list of reagents and volumes used is available in Table S2.

Single-color compensation controls for monoclonal antibody reagents were prepared per manufacturer recommendations using UltraComp eBeads Plus Compensation Beads (Invitrogen, cat. #01-3333-42) and 1μL of each experimental reagent and washed as above. Compensation controls for the viability dye were prepared using the ArC Amine Reactive Compensation Bead Kit (Invitrogen, cat. #A10346) and 1μL of dye. The MR1 tetramer compensation control was prepared using 1e6 adult PBMCs under identical staining conditions to the experimental reagent. Controls corresponding to reagents targeting extracellular antigens were subsequently fixed as above.

### Flow Cytometry Acquisition and Data Analysis

All samples were acquired on a 5L Cytek Aurora analytical flow cytometer. Unmixed FCS files were exported for analysis using FlowJo (v10.10) or R (v4.5.0). FCS files were imported, annotated, and clustered using the SOMnambulate package (v1.0.0.1) to exclude non-lymphocytes and dead/dying cells using viability dye and scatter parameters. Briefly, a self-organizing map was generated to classify cells according to shared marker expression and nodes within the minimal spanning tree were grouped using k-nearest neighbor clustering. Live lymphocytes clusters were reclustered (k = 20) using the discriminatory markers CD3, CD4, CD8α, CD8β, CD56, and TCRγδ to identify major phenotypic lymphocyte populations. CD8α^+^ αβ T cells (i.e. CD3^+^CD4^-^CD8α^+^CD8β^+/-^CD56^-^TCRγδ^-^ events) were subsequently reclustered (k = 20) using the remaining panel markers after trimming extreme events (i.e. the top/bottom 0.01%). UMAP dimensionality reduction was performed using the uwot package (v0.2.2). Hierarchical gating was performed in FlowJo (v10.10) using unmixed and ungated FCS files to confirm marker expression observed on derived cellular clusters.

### Immunostaining: Single Cell Sequencing

Samples were thawed in batches of 4 subjects and prepared as described above for stimulation and immunostaining except for the initial plating concentration (2e6 cells/well). IL-12/IL-18-treated and unstimulated cells were separately blocked and stained with LIVE/DEAD Fixable Scarlet Dye (Invitrogen, cat. #L34986), library-specific TotalSeq-C hashing reagents (BioLegend, see Table S2), barcoded and fluor-conjugated MR1 (Immudex, cat. #ZA08004DXG) and CD1d Dextramers (Immudex, cat. #CSS005), and an antibody cocktail consisting of barcoded TotalSeq-C reagents and a minimal, fluor-conjugated sorting panel with matched, compatible clones targeting sorting markers (Table S2). Donor-specific unstained cellular samples were processed in parallel but without the addition of any experimental reagents. Stained cells were washed four times with 2% Staining BuCer before centrifugation and supernatant aspiration. Cell pellets were stored on ice until sorting. Bead- and cell-based compensation controls were prepared as above but without a fixation step for surface markers. A full list of reagents and volumes used is available in Tables S3 and S4.

### Fluorescence-Activated Cell Sorting

Samples were sorted using a 5L Cytek Aurora CS system with an 85 μm nozzle. Unstained cellular samples were used to subtract a per-donor autofluorescence signature and to determine optimal scatter parameter voltages. Separately hashed stimulated and unstimulated cells were resuspended in cold R10 medium, pooled at a 2:1 ratio by volume, and filtered using a Cell Strainer-capped 5mL test tube (Falcon, cat. #352235). Pooled, donor-specific samples were sorted into 150μL cold R20 medium (RPMI-1640 with 20% FBS & 1X Pen/Strep) in 1.5mL Protein LoBind tubes (Eppendorf, cat. #022431081) using the ’purity’ sort mode to yield enriched cellular fractions.

### Single Cell Sequencing

Enriched cellular fractions were re-quantified using a viability dye exclusion assay and were pooled per-donor using the cell number and proportions shown in Fig 2A. Batches of 4 subjects and 8 hashed libraries underwent droplet encapsulation and cDNA synthesis using the GEM-X Single Cell Gene Expression and Immune Profiling kit (10X Genomics, cat. #1000699). UMI-labeled libraries consisting of 1) mRNA-derived cDNA, 2) TotalSeq-C and Dextramer feature barcodes and 3) V(D)J-amplified cDNA were prepared in parallel according to manufacturer instructions by the University of Rochester Medical Center’s Genomics Research Center. Library concentration was measured by qPCR before normalization, per-batch pooling, and loading onto one lane of a NovaSeq X flowcell (Illumina).

### GEX Assay Data Pre-processing

Raw sequence data were demultiplexed, processed, and aligned to the GRCh38-2024-A human genome reference assembly with the Cell Ranger multi pipeline (v9.0.0) using default parameters. Library-specific feature-barcode matrices corresponding to each batch were processed using Seurat (v5.0) to remove low-quality cells and doublets before merging and integration by library using Harmony (v0.1). Nearest neighbor graphs were generated for the Harmony-corrected PCA cell embeddings and were clustered using the smart local moving (SLM) algorithm with resolution 0.2-0.6. UMAP dimensionality reduction was performed in Seurat using the umap-learn python package. Per-cluster marker finding was performed (‘FindAllMarkers’) and each cluster was inspected for expression of canonical non-T cell marker transcripts (i.e. CD19 - B cells, CD14 - monocytes). Discrete B cell and monocyte clusters were excluded from downstream analyses and the filtered Seurat objects were saved for future loading as RDS objects.

### DSB Background Normalization

Both the raw and demultiplexed/filtered feature-barcode matrices were taken as input for batch-specific background normalization and denoising of the surface protein expression datasets using the DSB package (v2.0.0). RNA and protein library size metrics were derived by log-scaling per-cell library counts regardless of feature. "Empty" (i.e. non-cellular) droplet barcodes with intermediate RNA.size (1.0-2.5) and protein.size (1.0-3.0) values were isolated to construct a count profile of unbound protein features. Library normalization and denoising was performed using the background count matrix as well as isotype antibody profiles. Each normalized feature-barcode matrix was incorporated into the respective Seurat object as a new assay during initial QC and filtering.

### Batch Integration and Multimodal Analysis

Batch-specific RDS objects obtained from the above step (‘GEX Assay Data Pre-processing’) were re-processed with Seurat. Individual objects were merged and integrated by batch using Harmony to preserve potential biological variability across donors. Batch-corrected RNA and background-normalized protein assay data were used for weighted nearest neighbor (WNN) integrative analysis. Nearest neighbor graphs were again generated for the cell embedding graphs and were clustered using the smart local moving (SLM) clustering algorithms with resolution 0.2. Per-cluster marker finding was performed using both RNA and protein features and all surface CD4^+^ conventional T cell clusters that 1) lacked surface CD8α^+^ expression and 2) detectable ZBTB16 transcript were excluded from further analyses. CD8α^+^ αβ T cells were re-integrated and re-clustered using a clustering resolution of 0.4 and an otherwise identical approach for downstream analyses. Statistical testing for differential feature expression was performed using the Wilcoxon rank sum test as implemented in Seurat or ggpubr (v0.6.0) unless otherwise specified.

### Pseudobulk Analysis

Pseudobulk analyses were performed with the PCAtools (v2.21.0) package. Briefly, per-cluster pseudobulk profiles were derived using RNA and non-hashing protein features (‘AggregateExpression’) and principal components analysis was performed (‘pca’) before visualization.

### Regulon Analysis

Transcription factor activity inference analysis was performed with the decoupleR (v2.9.7), dorothea (v1.7.4) and SCpubr (v2.0.2) packages. Briefly, the dorothea human database was loaded and used to score each cell’s expression of transcription factors and their associated downstream targets.^56,84^ A weighted mean enrichment score for each regulon was derived (‘run_wmean’) and visualized per-cell type with SCpubr.

### TCR Analysis

Per-library filtered contig annotation files were loaded and combined into per-cell clones with scRepertoire (v2.3.4). Any cells with >2 TCR chain sequences were excluded from subsequent analyses. Clone size estimates using TCRα/β CDR3 amino acid sequences were performed relative to each cell population as shown in Figure 5A. Clonal overlap and diversity calculations were performed for each cell population by exporting per-cell metadata features from the final Seurat object and joining them to the TCR contig data frame. Relevant TCR metadata were exported to the final integrated Seurat object for further visualization.

### Functional Annotation

DiCerential gene expression was determined with Seurat (‘FindMarkers’) for specified groups of cells using standard significance (p_adj_ < 0.05) and fold change (|FC| > 1.0). Log_2_ fold change values were filtered and ordered for GSEA using clusterProfiler and the *Homo sapiens* Gene Ontology database (‘gseGO’). Single cell gene set variation analysis (GSVA) was performed with escape (v2.3.4) using the Human MSigDB Hallmark_Interferon_Gamma_Response gene set (M5913) and visualized with Seurat.

### Data Visualization & Statistical Testing

All data visualization was performed in R with ggplot2 (v3.5.2), SCpubr (v2.0.2), or Nebulosa (v3.21) unless otherwise specified. All statistical testing was performed with the ggpubr (v0.6.0) package unless otherwise specified.

### Data and Materials Availability

GEO submission of all raw and processed data is currently in process.

All scripts and parameters used for data processing and analysis will be made available via GitHub.

## Acknowledgements

Research reported in this publication was supported by the URMC MSTP (NIH MSTP grant: 5T32GM007356) and by the URMC Dept. of Pediatrics Small Grants Program.

Subfigure cartoons and final figure layouts were made in BioRender.

---

**Figure S1:**
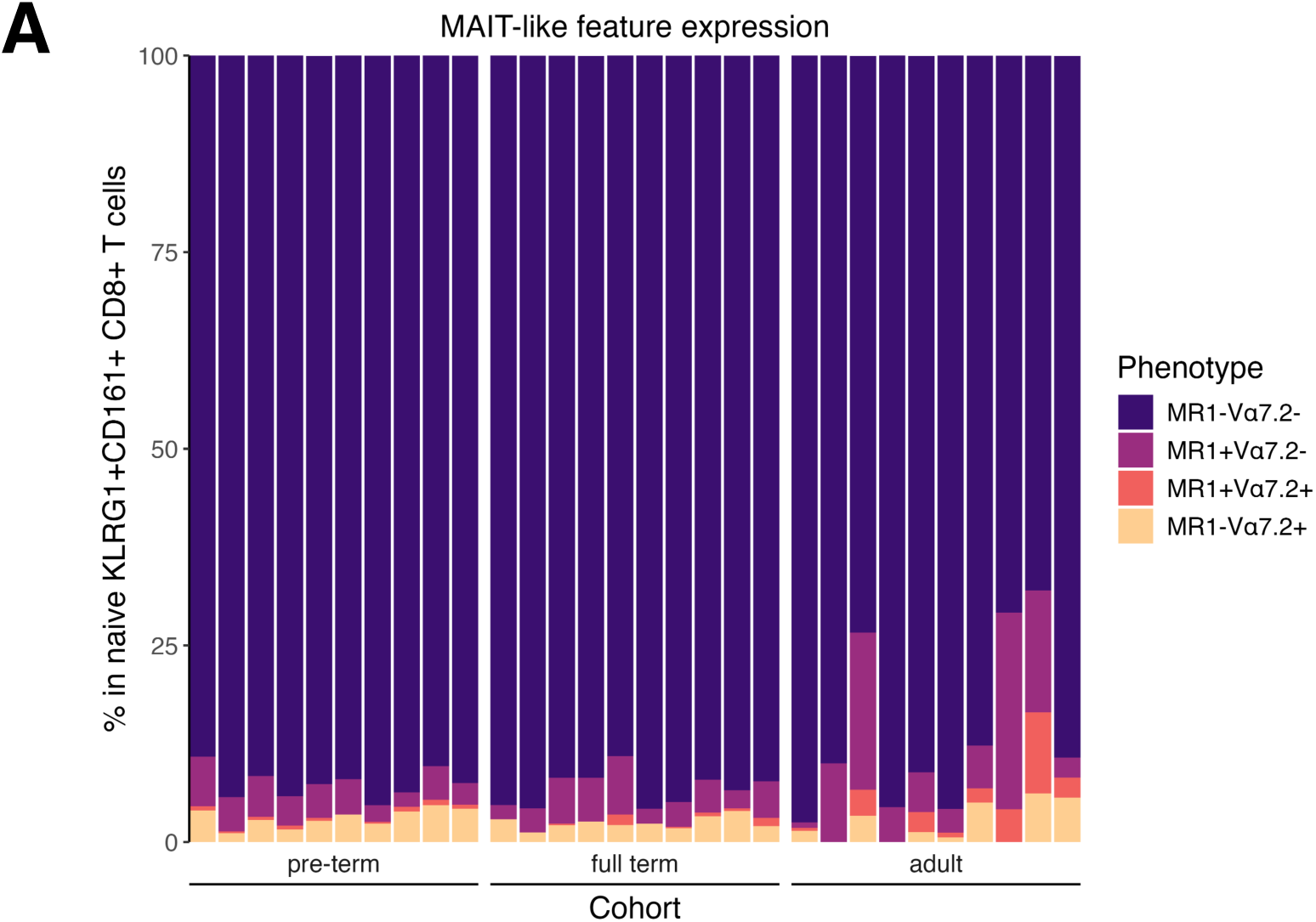
Infant naïve KLRG1^+^CD161^+^ CD8α^+^ αβ T cells minimally exhibit canonical MAIT features. A) Relative frequency of phenotypic populations defined by expression of the MAIT-associated features TCRVα7.2 and MR1:5-OP-RU tetramer binding among naïve KLRG1^+^CD161^+^ CD8^+^ αβ Τ cells. Infant and adult cohorts are grouped along the x-axis and ordered by increasing gestational or biological age.

**Figure S2:**
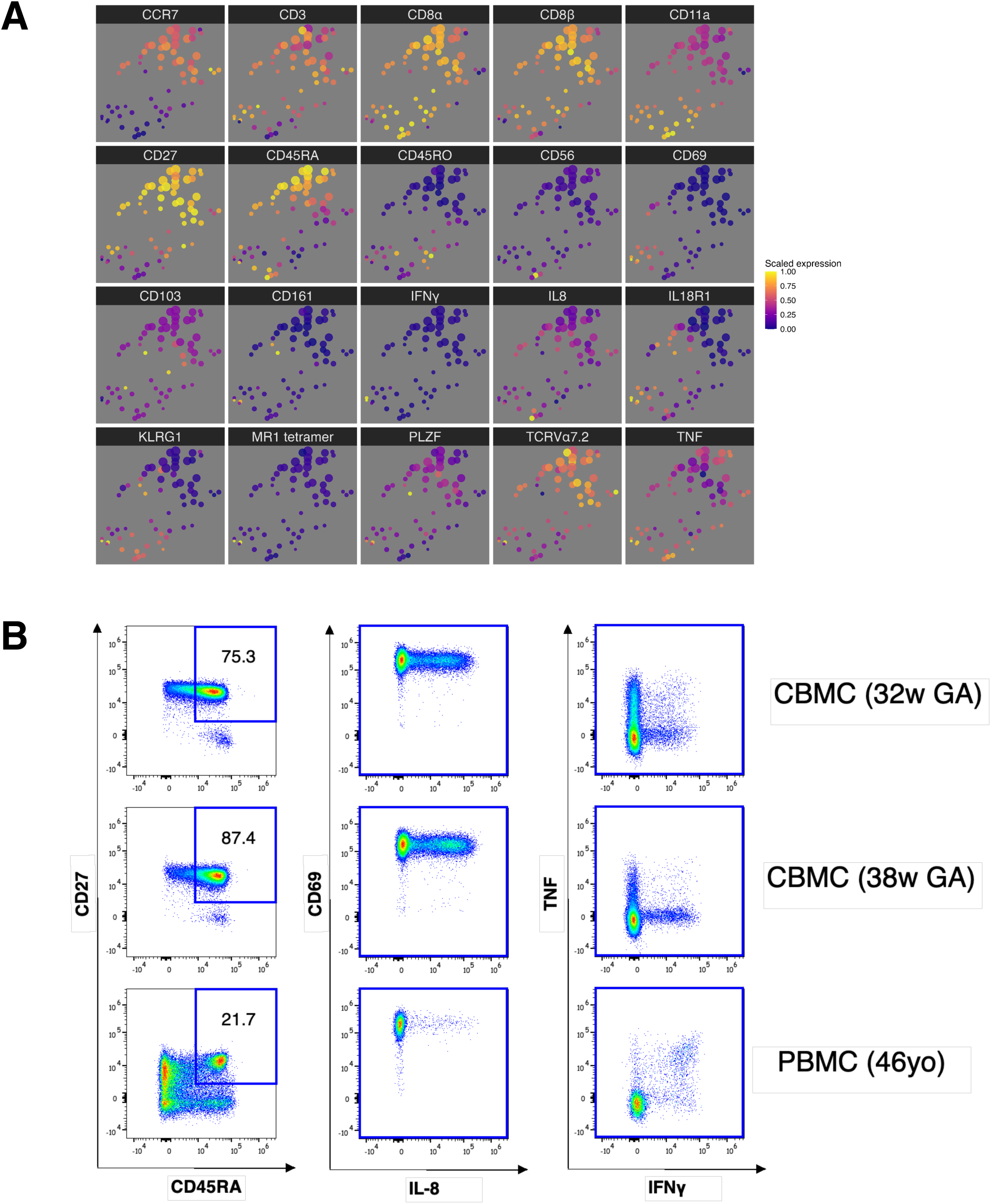
Fetal innate-like T cells are marked by co-expression of KLRG1 and CD161 and produce TNF following PMA/ionomycin stimulation. A) UMAP dimensionality reductions showing expression of all phenotypic features among FlowSOM-derived nodes. Expression values are scaled from 0 – 1 after trimming extreme events. B) Representative scatterplots of cytokine expression among naïve (CD45RA^+^CD27^+^) cellular populations following PMA/ionomycin stimulation.

**Figure S3:**
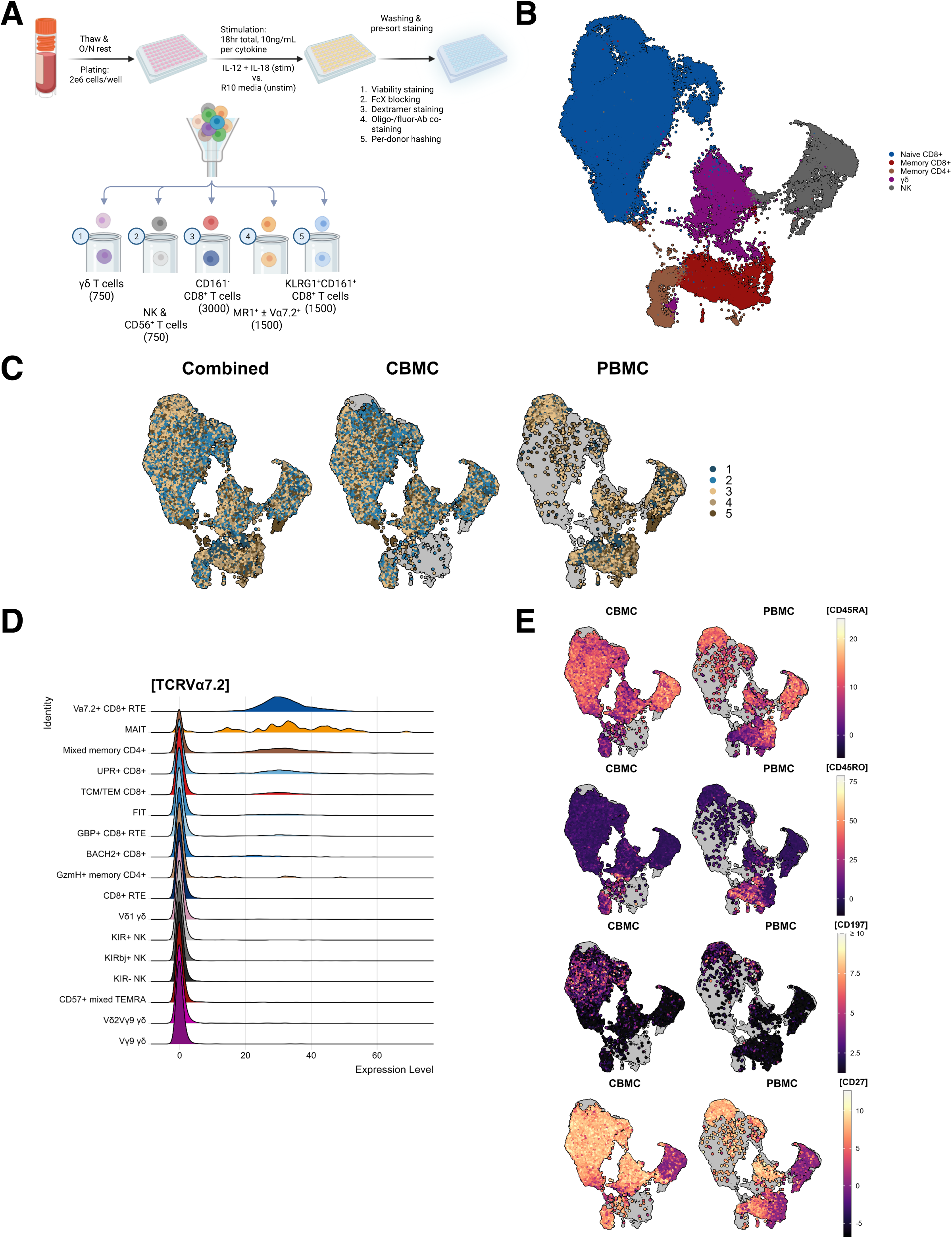
T and NK cells cluster by cell type and express memory phenotypic markers enabling their classification. A) Cartoon showing experimental workflow for CITE-seq labeling and cell sorting before droplet encapsulation. B) WNN UMAP with annotated cell types based on feature expression patterns as shown in Fig. 3C. Clusters are grouped by color i.e. naïve CD8α^+^ T (blue), memory CD8α^+^ T (red/orange), memory CD4^+^ αβ T (brown), γδ T (purple), or NK cells (grey). C) WNN UMAP split by age cohort and colored by batch. D) Ridge plot showing expression density of surface TCRVα7.2 as measured across specified lymphocyte clusters. E) Feature expression of major naïve/memory phenotypic markers among infant and adult cells.

**Figure S4:**
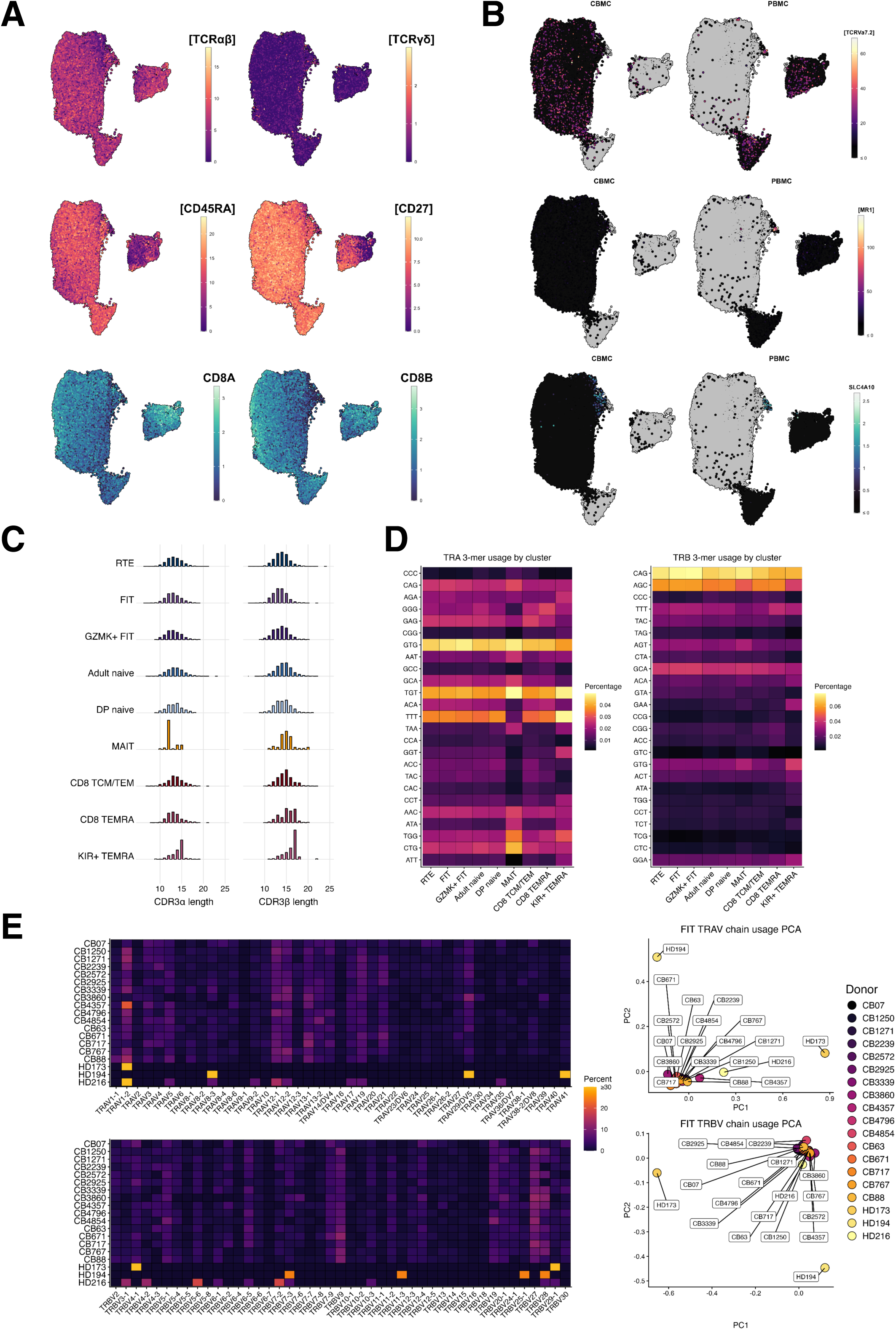
Conventional CD8α^+^ αβ T cells cluster by eWector/memory diWerentiation status and are clonally diverse while lacking canonical MAIT features. A) Feature plots showing expression of major phenotypic markers among strictly defined CD8α^+^ αβ T cells. B) Feature plots split by age cohort and showing expression of characteristic MAIT cell features. C) Calculated CDR3 lengths for TCRα (left) and TCRβ (right) chains among shown cell populations. D) 3-mer nucleotide usage frequencies for TCRα (left) and TCRβ (right) CDR3 regions among shown cell populations. E) Summary heatmaps showing relative frequency of TRAV (top) and TRBV (bottom) chains among FIT cells from the shown donors. Associated principal components analysis plots show the relative arrangement of per-donor FIT cells based on their TRAV (top) and TRBV (bottom) chain usage pattern.

**Figure S5:**
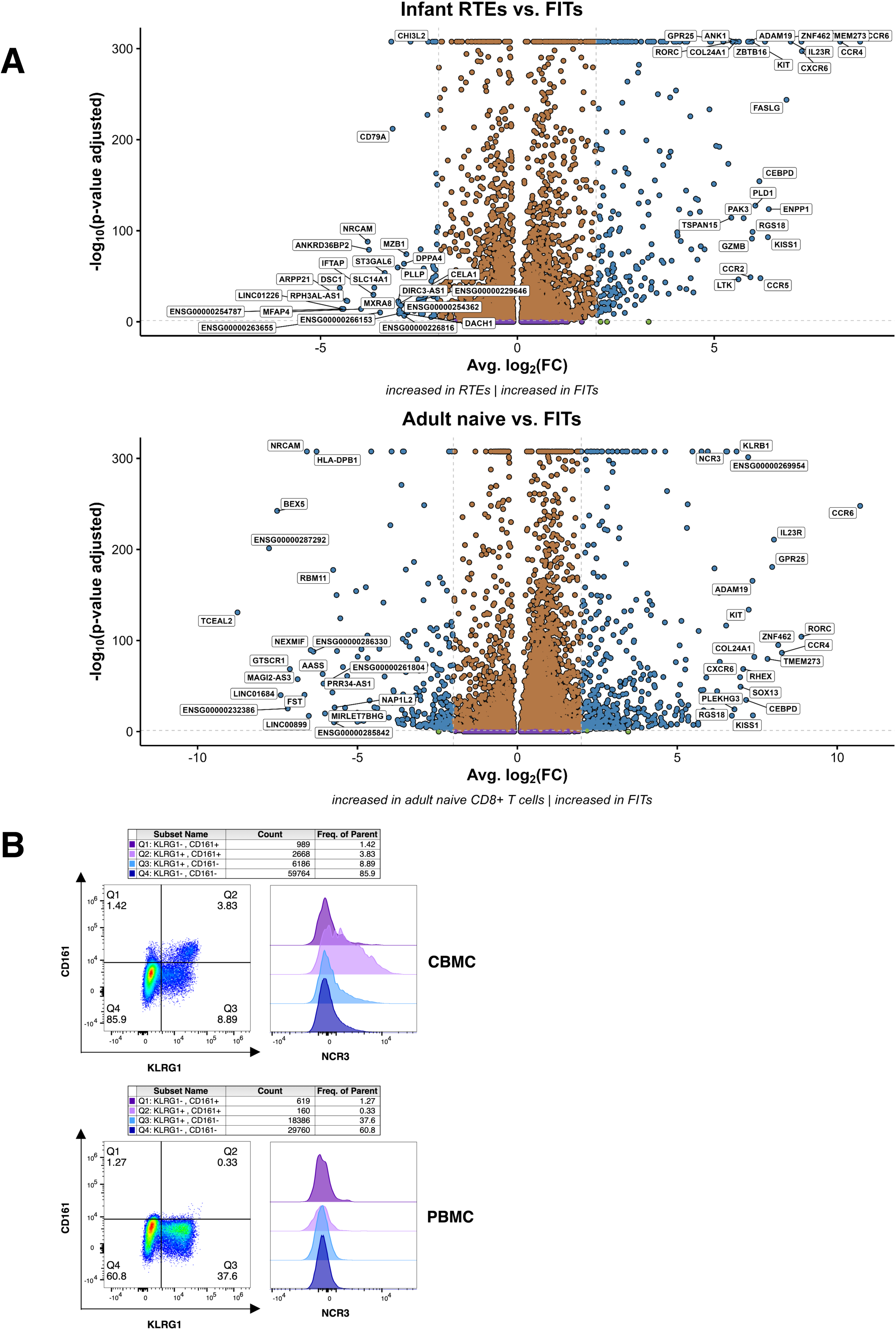
FITs are enriched for expression of type 17 immunity-related transcripts and NCR3 protein relative to other naïve CD8α^+^ αβ T cells. A) Volcano plots showing diqerentially expressed transcripts between FIT and infant RTE (top) and FIT and adult (bottom) naïve CD8α^+^ αβ T cell clusters. B) Representative CBMC and PBMC samples showing NCR3 expression among KLRG1/CD161 phenotypic subsets of naïve CD8α^+^ αβ T cells.

**Figure S6:**
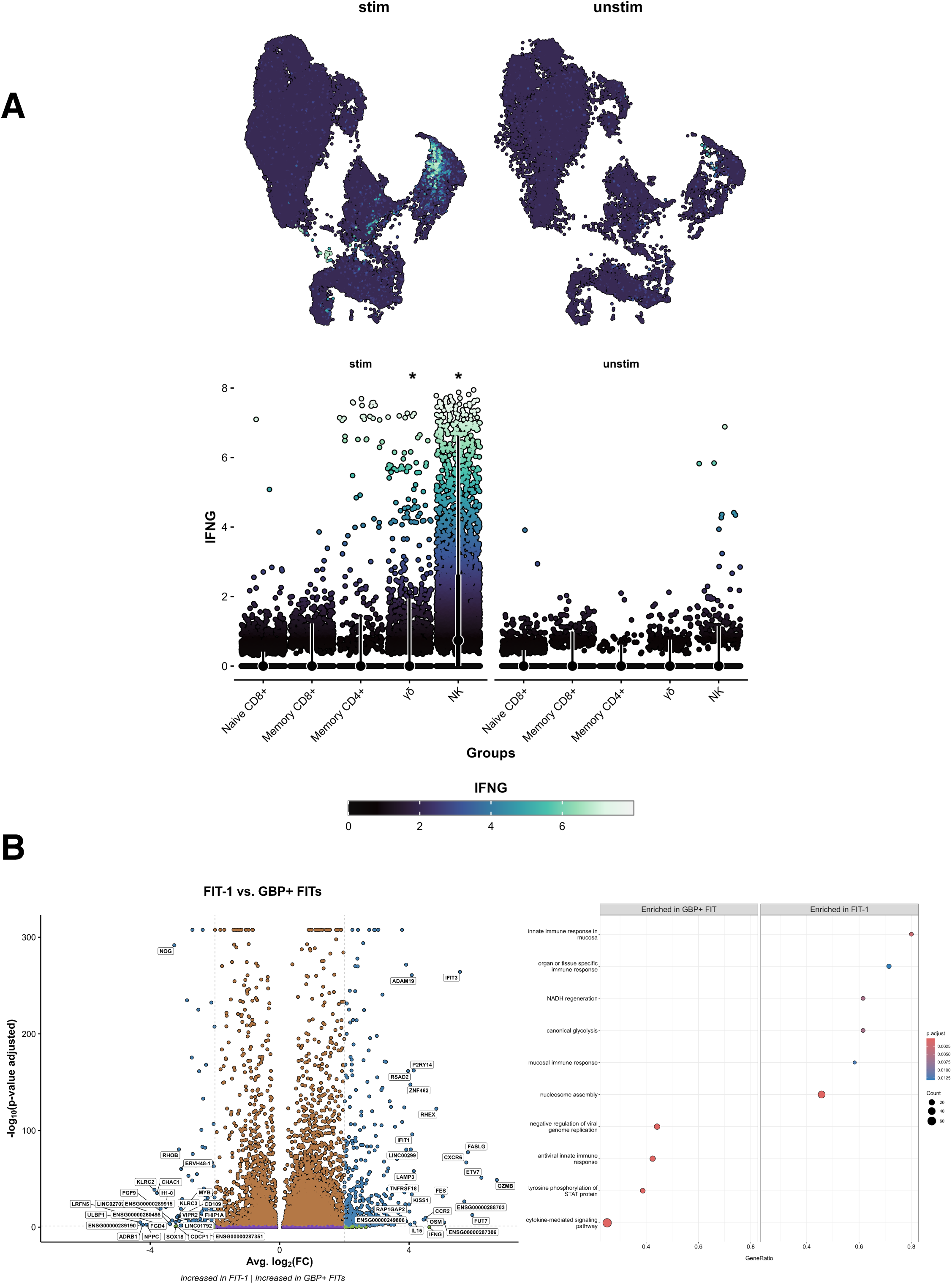
NK and γδ T cells, but not naïve CD8α^+^ αβ T cells, produce IFNγ following IL-12/IL-18 stimulation. A) Feature and geyser plots showing expression of IFNG transcript in unstimulated and stimulated cells. Significance (* - p < 0.001) for comparisons of feature expression level between stimulation conditions was determined with the Wilcoxon rank sum test. B) Volcano plot showing diqerentially expressed transcripts and GO GSEA significance plot derived from comparisons between between GBP+ FIT and FIT-1 naïve CD8α^+^ αβ T cell clusters.

**Table S1: Available donor metadata and batch processing details for single cell sequencing of CBMC/PBMC samples.**

**Table 2-2: Experimental reagents and volumes used for flow cytometric immunophenotyping of CBMC/PBMC samples.**

All volumes listed are in microliters except for those reagents which were diluted and used with predetermined volumes as recommended by the manufacturer. Compartment designations are as follows: ‘E’ – extracellular, ‘I’ – intracellular, ‘N’ – nuclear.

**Table 2-3: TotalSeq-C and Immudex experimental reagents and concentrations used for surface labeling of CBMC/PBMC samples before downstream CITE-seq profiling.**

**Table 2-4: Experimental reagents and volumes used for surface co-staining before cell sorting of CBMC/PBMC samples.**

## Notes

### Competing Interest Statement

The authors have declared no competing interest.

